# Selection for rapid uptake of scarce or fluctuating resource explains vulnerability of glycolysis to imbalance

**DOI:** 10.1101/2019.12.12.873315

**Authors:** Albertas Janulevicius, G. Sander van Doorn

## Abstract

Glycolysis is a conserved central pathway in energy metabolism that converts glucose to pyruvate with net production of two ATP molecules. Because ATP is produced only in the lower part of glycolysis (LG), preceded by an initial investment of ATP in the upper glycolysis (UG), achieving robust start-up of the pathway upon activation presents a challenge: a sudden increase in glucose concentration can throw a cell into a self-sustaining imbalanced state in which UG outpaces LG, glycolytic intermediates accumulate and the cell is unable to maintain high ATP concentration needed to support cellular functions. Such metabolic imbalance can result in “substrate-accelerated death”, a phenomenon observed in prokaryotes and eukaryotes when cells are exposed to an excess of substrate that previously limited growth. Here, we address why evolution has apparently not eliminated such a costly vulnerability and propose that it is a manifestation of an evolutionary trade-off, whereby the glycolysis pathway is adapted to quickly secure scarce or fluctuating resource at the expense of vulnerability in an environment with ample resource. To corroborate this idea, we perform evolutionary simulations of a simplified yeast glycolysis pathway consisting of UG, LG, phosphate transport between a vacuole and a cytosol, and a general ATP demand reaction. The pathway is evolved in constant or fluctuating resource environments by allowing mutations that affect the (maximum) reaction rate constants, reflecting changing expression levels of different glycolytic enzymes. We demonstrate that under limited constant resource, the population evolves to a genotype that is balanced but exhibits strongly imbalanced dynamics under ample resource conditions. Furthermore, when resource availability is fluctuating, the imbalanced phenotype enjoys a fitness advantage over balanced dynamics: when glucose is abundant, imbalanced pathways can quickly accumulate glycolytic intermediate FBP as intracellular storage that is used during periods of starvation to maintain high ATP concentration needed for growth. Our model further predicts that in environments with fluctuating resource, competition for glucose can result in stable coexistence of balanced and imbalanced cells, as well as repeated cycles of population crashes and recoveries that depend on such polymorphism. Overall, we demonstrate the importance of ecological and evolutionary arguments for understanding seemingly maladaptive aspects of cellular metabolism.

## Introduction

In many organisms, glycolysis is an essential pathway in energy metabolism that converts glucose to pyruvate with net production of two ATP molecules per glucose molecule^1^. Net formation of ATP occurs in the lower part of glycolysis (LG) which is preceded by initial investment of ATP in the upper part of glycolysis (UG). Such a “turbo design” of the pathway carries an inherent risk: a sudden increase in glucose levels can push the pathway into a self-sustaining imbalanced state, where UG outpaces LG, glycolytic intermediates accumulate and the cell is unable to maintain high ATP concentration needed to support cellular functions^2,3^. In yeast, such a phenotype is usually associated with mutants of the trehalose metabolism^2,4,5^. However, wild-type (WT) yeast cells are vulnerable as well: a switch of growth substrate from galactose to glucose renders 7% of cells non-viable^3^. This is an example of substrate-accelerated death, a wider phenomenon observed in prokaryotes and eukaryotes when cells are unable to grow when exposed to excess substrate that previously limited growth^6–8^.

The co-occurrence of a balanced and an imbalanced state in yeast glycolysis is well captured by a generalized core glycolysis model (Figure 1) developed previously by van Heerden et al.^3^ The model predicts two stable states, one yielding a steady-state concentration of the intermediate metabolite fructose-1,6-bisphosphate (FBP) and a high ATP concentration (balanced state), and the other characterized by steady accumulation of FBP and depletion of ATP and intracellular inorganic phosphate pools (imbalanced state). The key factor determining the fate of the system is the dynamics of inorganic phosphate (P_i_) during the start-up of glycolysis. According to the model, the transition to excess glucose proceeds as follows 3. Upon sudden glucose exposure, the rate of UG initially exceeds that of LG (*v*_up_ > *v*_lo_), causing FBP to increase. For a balanced steady state, *v*_lo_ has to accelerate and catch up with *v*_up_. This challenge is more difficult if UG activity (*v*_up_) is higher, e.g. due to a higher expression level of UG enzymes or higher glucose concentration. Accumulation of FBP binds P_i_ and will cause a drop in its concentration in the cytosol. Because FBP and P_i_ are both substrates for LG, the increase in FBP will tend to speed up *v*_lo_, whereas the associated decrease in P_i_ will tend to slow it down. Which of these two effects is dominant determines the trajectory of the system. If P_i_ concentration remains sufficiently high, *v*_lo_ will increase to become equal with *v*_up_, and a balanced steady state will be established. Otherwise, P_i_ will become a limiting factor for LG and *v*_lo_ will not accelerate fast enough to catch up with *v*_up_, causing the system to collapse into the imbalanced state. Once caught in the imbalanced state, cells are trapped, because P_i_ mobilized from the vacuole maintains the imbalance: at low concentrations of P_i_ and ATP, an imported P_i_ molecule enhances *v*_lo_, but the concomitant production of 2 ATP molecules increases *v*_up_ twice as much as *v*_lo_. Given that imbalanced cells exist in an alternative stable metabolic state, random initial variation in enzyme and metabolite concentrations can be enough to drive a subpopulation of cells into the imbalanced state. This explains why both balanced and imbalanced cells can be present in an isogenic population upon transition to excess glucose after starvation^3^.

**Figure 1.**
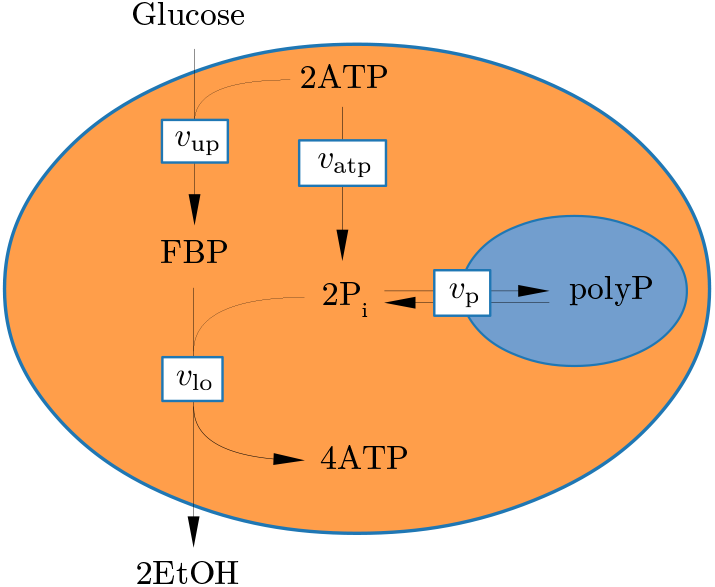
A generalized core model of glycolysis^3^ considers the intracellular concentrations of the glycolytic intermediate fructose-1,6-bisphosphate (FBP), ATP and inorganic phosphate (P_i_), and four reactions (arrows): (i) a lumped upper glycolysis reaction that produces FBP from extracellular glucose with rate *v*_up_, (ii) a lumped lower glycolysis reaction that generates ATP and the waste product ethanol (EtOH) at rate *v*_lo_, (iii) ATPase reaction reflecting general ATP demand in the cell at rate *v*_atp_, and (iv) reversible phosphate transport between the cytosol and the vacuole at rate *v*_p_.

van Heerden et al. suggests that vulnerability of glycolysis to imbalance arises from the fundamental design of the pathway and cannot be fully prevented by regulatory mechanisms^3^. However, the analysis of the full kinetic glycolysis model shows that quicker liberation of P_i_ by enhancing ATPase activity, activation of the glycerol formation branch, futile trehalose cycling, or quicker import of P_i_ into the cytosol from the vacuole can all markedly decrease the probability of reaching the imbalanced state^3^. Furthermore, the presence of trehalose cycling combined with experimentally observed trehalose-6-phosphate mediated inhibition of hexokinase^9^ can remove the existence of metabolic imbalance in the model altogether. It is therefore puzzling that 7% of WT yeast cells fall into the imbalanced state upon a sudden increase in glucose availability. Why have WT cells not evolved such mechanisms to completely eliminate the risk of imbalance? One possible evolutionary explanation is that although imbalanced *v*_up_ and *v*_lo_ are dangerous to the cell, regulatory mechanisms to keep them tightly balanced, or constitutive higher expression of LG enzymes are just too costly relative to the fitness benefit of avoiding substrate-accelerated death. Yet, given the potential of a 7% increase in survival, these costs must be assumed to be substantial. An alternative hypothesis that we propose here is that imbalanced *v*_up_ and *v*_lo_ are not always detrimental to the cell, but may, in fact, be adaptive under a range of natural conditions. In particular, allocating a larger fraction of enzymatic capacity to UG at the expense of LG would allow cells to acquire glucose from the environment faster, increasing their competitive advantage under conditions of low resource availability. Moreover, in variable environments, glucose may normally run out before metabolic imbalance becomes irreversible, so that periods of starvation would restore normal levels of glycolytic intermediates and cells would be protected from substrate-accelerated death. From this perspective, the vulnerability of cells to fall into the imbalanced state in rich and constant environments (e.g., typical lab conditions) can be interpreted as the result of an evolutionary trade-off: adaptations of the glycolysis pathway that improve its performance under conditions of low or varying glucose make it vulnerable to imbalance at constant high glucose concentrations. In other words, we suggest that imbalanced dynamics in WT yeast cells are observed because cells are adapted to a different glucose availability regime than the one used in the experiments^3^.

To investigate this idea, we performed evolutionary simulations of a population of yeast cells with the simplified yeast glycolysis pathway shown in Figure 1, subject to different glucose availability regimes. Variation in the population was introduced by mutating (maximum) reaction rate constants of the pathway, reflecting changing expression levels of the glycolytic enzymes. Cells contributed to future generations in proportion to their growth rate, so that natural selection acted on the simulated populations to improve the functionality of the pathway in the current environment. We then quantified the likelihood of cells with the evolved pathways to fall into the imbalanced state upon transitioning to excess glucose. Our results demonstrate that the regime of glucose availability that cells have previously adapted to has a marked effect on their measure of balancedness. The model also predicts a range of environmental conditions where balanced and imbalanced cells can stably coexist in the population, and where such polymorphism drives periodic crashes and recoveries of the population. We discuss these results in relation to the tragedy of the commons and evolutionary suicide to illustrate how eco-evolutionary mechanisms can shed new light on seemingly maladaptive aspects of cellular metabolism.

## Model and methods

### Model description

We model a population of yeast cells that metabolize glucose in a chemostat. To model glycolysis, we employ a generalized core model^3^ comprising four reactions (Figure 1), with the addition of explicit glucose dynamics and phosphate depletion from the yeast vacuole:

i. Upper glycolysis is modeled to exhibit irreversible two-substrate Michaelis-Menten kinetics. Phosphofructokinase, an enzyme of upper glycolysis, is allosterically inhibited by ATP^1,10,11^, hence the reaction rate of upper glycolysis *v*_up_ contains an inhibition term with the inhibitor constant *K*_i,atp_ for ATP:

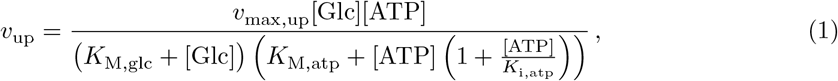

where *v*_max,up_ is maximal upper glycolysis rate, *K*_M,glc_ and *K*_M,atp_ are the Michaelis constants for glucose and ATP, respectively (similar notations are used below for corresponding parameters in other reactions).
ii. Lower glycolysis is assumed to follow irreversible three-substrate Michaelis-Menten kinetics. Its rate, *v*_lo_, is given by

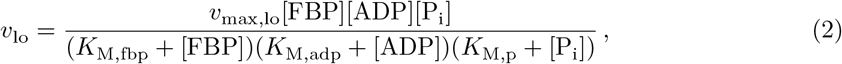

where [ADP] is found from a conserved quantity *a*_tot_ = [ATP] + [ADP].
iii. ATP consumption by all kinds of cellular processes (ATP demand) is modeled by a general ATPase reaction that follows first-order reaction kinetics with rate

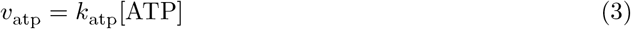

and reaction rate constant *k*_atp_.
iv. Phosphate is transported between the vacuole and the cytosol at rate

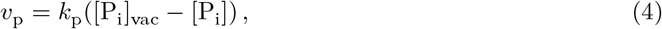

where [P_i_] and [P_i_]_vac_ are the phosphate concentration in the cytosol and the vacuole, respectively. When [P_i_] < [P_i_]_vac_, phosphate is transported from the vacuole into the cytosol (*v*_p_ > 0), and in the opposite direction when [P_i_] > [P_i_]_vac_ (*v*_p_ < 0). It has been observed that glycolytic intermediates accumulate in cells that undergo imbalanced dynamics until all phosphate from the vacuole is depleted^3,4,9^. We model the depletion of phosphate from the vacuole by assuming that [P_i_]_vac_ drops as the total concentration of phosphate imported into the cytosol [P_tot_] = [P_i_] + 2[FBP] + [ATP] increases:

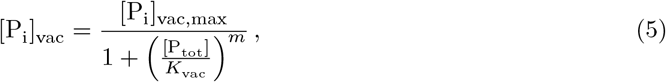

where [P_i_]_vac,max_ is the phosphate concentration in the vacuole when no phosphate in the cytosol is present, *K*_vac_ is the total concentration of phosphate in the cytosol that reduces [P_i_]_vac_ to one half of [P_i_]_vac,max_, and *m* > 0 determines whether phosphate depletion sets in gradually (small *m*) or suddenly (large *m*).

Metabolite concentrations in the cell are affected not only by glycolysis reactions, but also by dilution due to the increase in volume *V* of a growing cell. The decrease in metabolite concentration *c* due to dilution can be found from the conservation of the amount of metabolite *cV* in the cell as its volume increases:

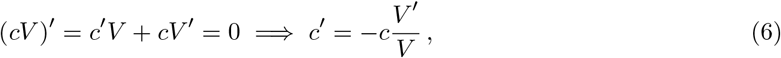

where the derivatives (denoted by the prime symbol) are with respect to time.

The dynamics of metabolites that participate in glycolysis reactions in a growing cell are thus governed by the following ordinary differential equations:

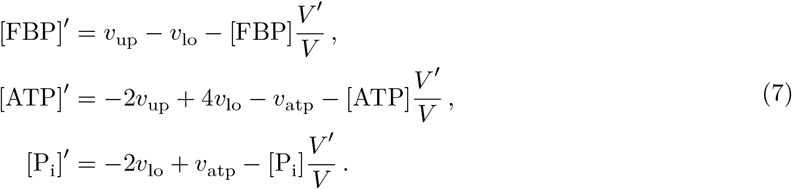

*v*_max,up_, *v*_max,lo_, *k*_atp_ and *k*_p_, the parameters of the metabolic pathway, reflect the expression levels of respective enzymes and define the genotype of the model cell. Since we aim to study evolutionary adaptation of this metabolic network, we must next specify its connection to growth and survival, the two key components of cellular fitness. In our model, glycolysis is coupled to fitness by the general ATPase reaction. We assume that the flux through this reaction (*v*_atp_) is first allocated to cover cellular maintenance costs (*v*_atp,c_) and that any remaining flux (*v*_atp,g_) is invested in cell growth, i.e. the production of new cell biomass, leading to an increase in cell volume *V*. The maintenance costs are further decomposed into *v*_atp,e_, the ATP demand required for expressing the glycolytic enzymes, which therefore may vary between cell genotypes, and the ATP required by other transcription and general cell maintenance processes (*v*_atp,m_), which is assumed to be equal between genotypes. Hence,

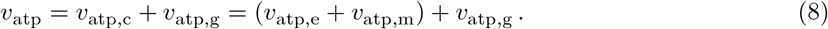

Because the cell maintenance flux *v*_atp,c_ is an obligatory component of the energy budget, cells are faced with an energy deficit when *v*_atp_ < *v*_atp,c_ (or, equivalently, when *v*_atp,g_ < 0). This occurs at low intracellular ATP concentration during periods of starvation or glycolytic imbalance. We assume that cells can buffer short periods of negative energy balance by drawing on internal reserves, but that they eventually die when starvation or imbalance persists. To model the deteriorating condition of a starving cell, we introduce a variable *H* that reflects cell health. Cell health decreases when ATP production falls short of meeting the energy demands for maintenance (i.e., when *v*_atp,g_ < 0). A cell dies when *H* decreases to *H*_min_ = 0, but, if starvation ends before this point is reached, the cell can recover. In fact, when *v*_atp,g_ becomes positive after a period of starvation, it is first invested into replenishing the internal reserves (modeled as an increase in *H*). When a cell is at its maximum health *H*_max_, and *v*_atp,g_ > 0, the cell will increase in volume. We assume that the amount of ATP needed to produce a unit of new cell volume is constant and independent of cell volume or genotype. In other words, the increase in cell volume is proportional to the total amount of ATP converted by the flux *v*_atp,g_ in the cell,

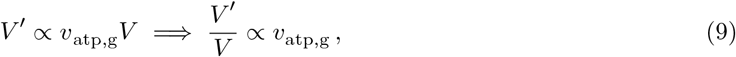

i.e. the rate of increase in the fractional cell volume is proportional to *v*_atp,g_. This will yield an exponential increase in cell volume at constant *v*_atp,g_, which is consistent with experimental measurements of yeast cell growth^12^.

To find the proportionality constant, we assume that a cell with balanced glycolysis in an environment with constant 2 mM glucose and the reference genotype reported by Heerden et al.^3^ (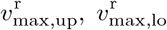, 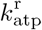 and 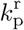) will double its volume in time *τ*_g_. Since under these conditions the ATP demand of the core glycolysis pathway is 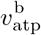, the expression cost of the reference genotype 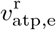 is chosen to yield a positive reference growth flux 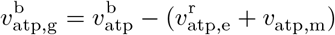 (Table 1). The dynamics of cell growth in our model is therefore

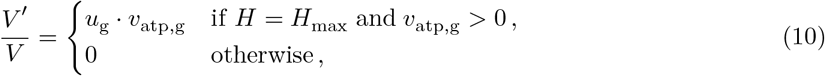

where

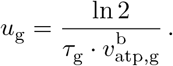

**Table 1.** Parameter values used in the simulations.

| Parameter | Value | Notes |
| --- | --- | --- |
| Extended core glycolysis model |  |  |
| $v_{\text{max,up}}^r$ | 10 mM · min<sup>-1</sup> | Ref. 3 |
| $K_{\text{M,glc}}$ | 0.1 mM | Chosen to be approximately equal to $K_{\text{M,glc}}$ of hexokinase<sup>11</sup> |
| $K_{\text{M,atp}}$ | 0.1 mM | Ref. 3 |
| $K_{\text{i,atp}}$ | 3 mM | Ref. 3 |
| $a_{\text{tot}}$ | 5 mM | Ref. 3 |
| $v_{\text{max,lo}}^r$ | 10 mM · min<sup>-1</sup> | Ref. 3 |
| $K_{\text{M,fbp}}$ | 1 mM | Ref. 3 |
| $K_{M,adp}$ | 0.1 mM | Ref. 3 |
| $K_{M,p}$ | 2 mM | Ref. 3 |
| $k_{atp}^r$ | $10 \text{ min}^{-1}$ | Ref. 3 |
| $k_p^r$ | $0.3 \text{ min}^{-1}$ | Ref. 3 |
| $[P_i]_{vac,max}$ | 10 mM | Ref. 3 |
| $K_{vac}$ | 250 mM | |
| $m$ | 4 | |
| $[FBP]_0$ | 2 mM | Ref. 3 |
| $[ATP]_0$ | 1 mM | Ref. 3 |
| $[P_i]_0$ | 10.4 mM | Ref. 3, used in all simulations except to determine $B_g$ |
| | 10 mM | Value used to determine $B_g$ |
| Cell |  |  |
| $\sigma$ | 0.1 | |
| $\mu$ | $10^{-2}$ | |
| $V_c$ | 3.35 fL | Cytosol volume of a spherical cell of diameter 2 $\mu\text{m}$ with 20% volume of organelles<sup>18</sup> |
| $\tau_g$ | 90 min | Ref. 19 |
| $\tau_d$ | 420 min | Estimated from Ref. 3 |
| $v_{atp}^b$ | $12.7 \text{ mM} \cdot \text{min}^{-1}$ | Estimated from Ref. 3 |
| $v_{atp}^i$ | $0.46 \text{ mM} \cdot \text{min}^{-1}$ | Estimated from Ref. 3 |
| $v_{atp,e}^r$ | $5 \text{ mM} \cdot \text{min}^{-1}$ | |
| $v_{atp,m}$ | $0 \text{ mM} \cdot \text{min}^{-1}$ | |
| Simulation |  |  |
| $N_0$ | 50 000 | Chemostat |
|  | 10 000 | NCG scenario |
|  | 12 000 | Two-genotype coexistence |
| $N^*$ | 10 000 | |
| $N_{tr}$ | 100 | |
| $V_{ch,0}$ | 10 nL | |
| $d_0$ | $1 \times 10^{-6} \text{ min}^{-1}$ | |
| $D$ | $5 \text{ min}^{-1}$ | NCG scenario |
| $t_s$ | 0 min | |
| $t_{ms}$ | 10 000 min | |
| $t_{me}$ | 500 000 min | |
| $t_e$ | 800 000 min | |
| $\Delta t_p$ | 5 min | |
| $\Delta t_s$ | 1 min | |
| ODE solver |  |  |
| $atol_c$ | $1 \times 10^{-5} \text{ mM}$ | Absolute tolerance for metabolite concentration |
| $rtol_c$ | $10^{-5}$ | Relative tolerance for metabolite concentration |
| $atol_v$ | 0.01 fL | Absolute tolerance for cell volume |
| $\text{rtol}_v$ | 0 | Relative tolerance for cell volume |
| $\text{atol}_h$ | $10^{-2}$ | Absolute tolerance for cell health |
| $\text{rtol}_h$ | 0 | Relative tolerance for cell health |

Cell health dynamics is similarly scaled by assuming that a cell with the reference genotype in an imbalanced state will die in time *τ*_d_. Under these conditions, a cell has a small ATPase flux 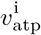 and thus a negative reference growth flux 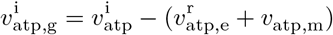. Cell health dynamics is therefore

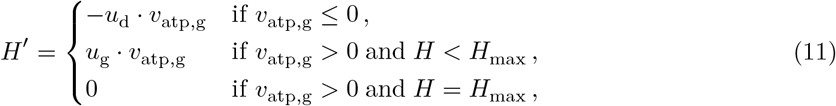

where

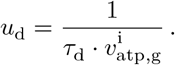

Given that parameters *v*_max,up_, *v*_max,lo_, *k*_atp_ and *k*_p_, which constitute the genotype of the cell, are proportional to the expression levels of glycolytic enzymes, we utilize these parameters to quantify the cost of expression, *v*_atp,e_. Because the expression costs of enzymes is difficult to estimate or measure experimentally^13^, we chose to investigate two cost functions,

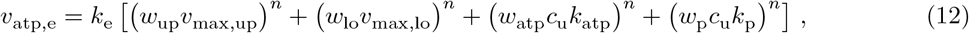

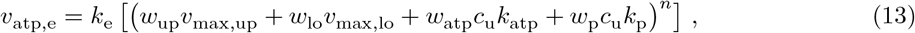

where *c*_u_ = 1 mM is the unit concentration, introduced for dimensional consistency, *w*_up_, *w*_lo_, *w*_atp_, *w*_p_ are weights of respective parameters on the total cost and *k*_e_ is a normalizing factor that assigns expression cost 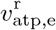 to the reference genotype. Unless indicated otherwise, Equation 12 is used with *w*_up_ = *w*_lo_ = *w*_atp_ = *w*_p_ = 1 and *n* = 4, and Equation 13 is referred to as the alternative cost function. The rationale to consider similar weights for multi-step, multi-enzyme pathways of UG and LG, and single-protein ATP demand and phosphate transport reactions stems from the fact that a multi-step pathway can be sped up by increasing the rate of one rate-limiting reaction (e.g., hexokinase or phosphofructokinase in UG, or pyruvate kinase in LG^1,14^). The nonlinearity in the cost function ensures that glucose flux through the pathway, and therefore ATP production, cannot be increased infinitely by the cell by increasing the total level of glycolytic enzyme expression.

Once the cell volume has increased to twice the standard cell volume *V*_c_, the cell divides. To prevent clonal subpopulations from dividing or dying synchronously, we introduce individual variability in the initial cell volume and the parameter *H*_max_. At the beginning of a simulation and after a cell division, each new (daughter) cell is assigned an individual uniformly distributed random value *H*_max_ ~ *U*(0.9, 1.1); a new (daughter) cell always starts with *H* = *H*_max_. Similarly, each new cell at the beginning of a simulation starts with initial uniformly distributed *V* ~ *U*(0.5*V*_c_, 1.5*V*_c_), whereas after cell division, only one of the daughter cells is assigned such a random volume, while the other daughter cell is left with the volume complementary to 2*V*_c_, i.e., the volume of the parent cell is divided between the two daughter cells. Daughter cells may be exposed to mutations of the genotype, which implies changing expression levels of corresponding glycolytic enzymes. Upon cell division, each parameter in the genotype of a daughter cell is modified with probability *μ*, or otherwise inherited from the mother cell. The modified value is drawn from a log-normal distribution

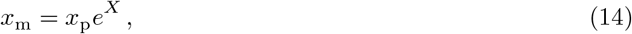

where *x*_p_ is the parental value, *x*_m_ is the mutated value in the daughter genotype and *X* ~ *N* (0, *σ*^2^) is a normally distributed random number with zero mean and standard deviation *σ*.

The final component of the model concerns the interaction between cells and the environment. A straightforward approach is to assume that the population of cells take up glucose, grow and divide in a chemostat chamber^15^. The glucose concentration in the chamber then changes due to uptake by cells, inflow and outflow of the medium, such that

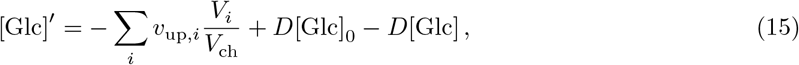

where the sum is over all cells in the population, *v*_up,*i*_ is the upper glycolysis rate of cell *i*, *V*_*i*_ is the volume of cell *i*, *V*_ch_ is volume of the chemostat chamber, [Glc]_0_ is the glucose concentration in the inflow medium and *D* is the dilution rate of the chemostat which is equal to *F*/*V*_ch_, where *F* is the medium flow rate. Cells are washed out from the chamber at a rate 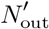 proportional to cell population size *N*,

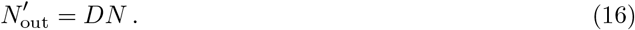

A chemostat is suitable to study a population of cells that compete for nutrient, because cells take up the nutrient and thus affect its concentration in the growth chamber. A mathematical analysis of the chemostat model shows that the nutrient concentration and the population size at steady-state depend on the maximum reproduction rate of cells^15^. Cells that reproduce faster, take up the nutrient faster, and thus, at steady-state, have a larger population size and leave less nutrient in the growth chamber. As nutrient uptake and reproduction rates of cells are evolving during evolutionary simulations, the steady-state nutrient concentration will also shift, making it difficult to determine the optimal evolutionary response of the metabolic network to a particular glucose availability regime. Therefore, we also considered an alternative model, where cell density is assumed to be so low that the consumption of glucose by cells has no noticeable effect on the glucose concentration in the chamber. In this version of the model, cells do not compete for glucose, but are limited by another resource, e.g., space in a biofilm that is attached to the wall of the chamber^16^. We refer to such conditions as the NCG (No Competition for Glucose) scenario. The NCG conditions can arise as a limiting case of the chemostat model where cells are attached to the substratum in a large chamber with a high flow rate, i.e. where *V*_ch_ → ∞ and *F* → ∞, while *D* = *F*/*V*_ch_ remains finite. The glucose uptake term in Equation 15 then vanishes and cells no longer affect glucose concentration in the chamber, i.e. [Glc] = [Glc]_0_. However, glucose concentration in the chamber is still affected by the medium inflow and outflow, allowing us to impose a particular glucose dynamics regime by adjusting *D* and [Glc]_0_. In this alternative model, cell loss rate from the attached biofilm in the chamber is

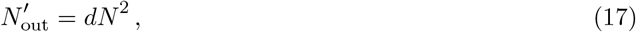

where *d* is the removal rate constant.

### Simulation procedure

The system of differential equations defined by Equations 7, 10 and 11 for each cell, and by Equation 15 for the glucose dynamics in the chemostat chamber, is solved in intervals of Δ*t*_p_ to obtain [FBP], [ATP], [P_i_], *V* and *H* dynamics for each cell and [Glc] dynamics in the chemostat chamber. Integration was carried out with the Dormand-Prince fifth-order Runge-Kutta method^17^ modified with non-negativity constraint for metabolite concentrations, i.e., if at the end of the integration step metabolite concentration *c* satisfied (atol_c_ + |*c*|rtol_c_) < *c* < 0, it was projected to zero. If after this any other *c* < 0 remained, the integration step was rejected and retried with a smaller step size. Between the integration intervals Δ*t*_p_, new cells are added to the population due to cell division, and cells are removed due to cell death and outflow from the chemostat.

A simulation is started with a population of *N*_0_ cells. To ensure that initial cell genotypes are sufficiently fit to survive and reproduce in a given glucose regime, initial values of the evolving parameters of each cell are drawn from a uniform distribution *U*(0.1*P*, 10*P*), where *P* is 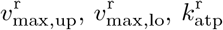 or 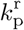 respectively.

Simulation time was divided into three segments: (i) from simulation start time *t*_s_ to mutation start time *t*_ms_ the mutation process was disabled, allowing the establishment of a viable steady-state population from the genetic variation created at the start of the run; this was necessary because many initial random genotypes were not viable under a given glucose availability regime; (ii) from *t*_ms_ to mutation end time *t*_me_ cells were exposed to mutations enabling a gradual evolution of the metabolic network, (iii) from *t*_me_ to simulation end time *t*_e_ the mutation process was disabled once again to allow only the fittest genotypes to remain in the population. As the speed of evolution is expected to depend on the population size, we sought to normalize the equilibrium population size at the beginning of segment (ii) to be approximately *N*^*^ across all simulations. To achieve that, we first performed a pre-simulation without mutation of the same duration as segment (i) with provisional values that regulate population size, i.e. *V*_ch,0_ for the standard chemostat model or *d*_0_ for the NCG scenario. After determining the steady-state size population size *N*_p_, full simulations with adjusted parameters 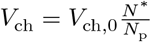 or 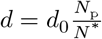 were run.

### Data analysis

Throughout the simulation, we tracked and saved the metabolite, volume and health dynamics of a random subpopulation of *N*_tr_ cells at time intervals of Δ*t*_s_. From this data, we find the fractional volume increase rate of tracked cells, *V*′/*V*, which is equivalent to cell reproduction rate *r* in population dynamics models. We also define an indicator to quantify the balancedness of the dynamics of the model core glycolysis pathway in a fluctuating environment. In a balanced cell, high external glucose coincides with high intracellular ATP, whereas in an imbalanced cell, high external glucose coincides with low intracellular ATP. The phenotypic balancedness of a cell, *B*_p,cov_ is therefore defined as covariance between the external glucose concentration and intracellular ATP concentration during an integral number of cycles in a periodic environment

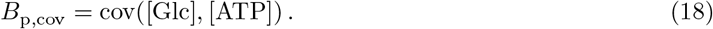

*B*_p,cov_ will have positive values if the dynamics of glycolysis is balanced and negative values if it is imbalanced. This measure is appropriate if glucose and ATP values oscillate regularly around their means; it is more difficult to interpret when the dynamics is irregular, as in the case of catastrophic dynamics (see Section *Evolution of increased imbalancedness…* below).

Under the NCG scenario studied here, the external glucose concentration in the chamber changes abruptly between a high value during the ON phase and a low value during the OFF phase because of a high *D* value. Therefore, a simpler and more easily interpretable measure of phenotypic balancedness, *B*_p,phs_, can be used by comparing the average ATP concentrations in the cell during ON and OFF phases:

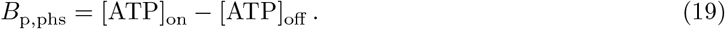

Also here, positive values indicate balanced dynamics (more ATP is produced and a cell grows faster during the ON phase), whereas negative values indicate imbalanced dynamics (more ATP is produced and a cell grows faster during the OFF phase).

Balanced and imbalanced glycolysis can exist as alternative steady-states for the same genotype. Therefore, we define the balancedness of a genotype (*B*_g_) as its propensity to exhibit balanced dynamics. Genotypic balancedness is determined by the following procedure, in part similar to the one described by van Heerden at al.^3^ For each genotype, we generate 100 random initial metabolite concentrations, apply a particular glucose concentration and simulate the metabolite dynamics for 300 min. The initial metabolite concentrations are normally distributed with either realistic constant means for all cells, [FBP]_0_, [ATP]_0_, [P_i_]_0_ (*B*_g,1_) or, in case of constant external glucose in NCG scenario, also the actual metabolite concentrations in evolved cells (*B*_g,2_), and realistic variation, CV = 6 %. We repeat the procedure for a range of glucose values, 2.00 mM, 1.95 mM, …, 0.05 mM. *B*_g_ is then the largest concentration of glucose that results in a balanced metabolism for all 100 random initial metabolite concentrations, or 0 mM otherwise. Thus, higher *B*_g_ indicates a more balanced genotype. To determine whether the metabolism is balanced or not in each of these simulations, we apply the following criterion. In a balanced phenotype under constant [Glc], [FBP] reaches a steady-state value. Equation 7 shows that at this steady-state

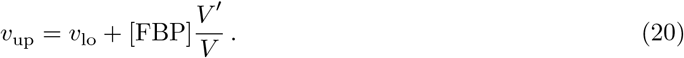

Taking into account that the cell needs time to reach metabolic steady state, the phenotype is considered balanced if Equation 20 holds true within 0.1% relative error for more than 10% of the simulation time.

## Results

We first investigate the evolution of the core glycolysis pathway in an environment with a constant concentration of glucose (NCG scenario, see *Model and methods*). In each of these simulations, the pathway evolves to optimize its performance in the particular glucose availability regime that is imposed externally. Next, we consider the evolution of the pathway in populations subject to competition for glucose in a chemostat, where adaptation of the pathway alters the ecological conditions experienced by the population. Due to this eco-evolutionary feedback, no single strategy may be optimal in a given environment, creating the possibility of more complex eco-evolutionary dynamics.

### Pathway adapted to scarce glucose exhibits imbalanced dynamics under ample glucose

Under NCG conditions, the core glycolysis pathway adapts to different constant levels of glucose availability by optimizing the expression levels of glycolytic enzymes (Figure 2A). The evolved expression pattern optimizes the balance between three selective forces. One component of selection favors an increase in *v*_max,up_, *v*_max,lo_ and *k*_atp_, because the increasing flux of glucose through the pathway enhances ATP production and cell growth rate. Next, there is a pressure to lower the genotype parameter values *v*_max,up_, *v*_max,lo_ and *k*_atp_ in order to reduce the cost of expression of the corresponding glycolytic enzymes. Finally, selection acts against *v*_max,up_ becoming too large compared to *v*_max,lo_ to avoid the loss of fitness due to cells falling into the imbalanced state.

**Figure 2.**
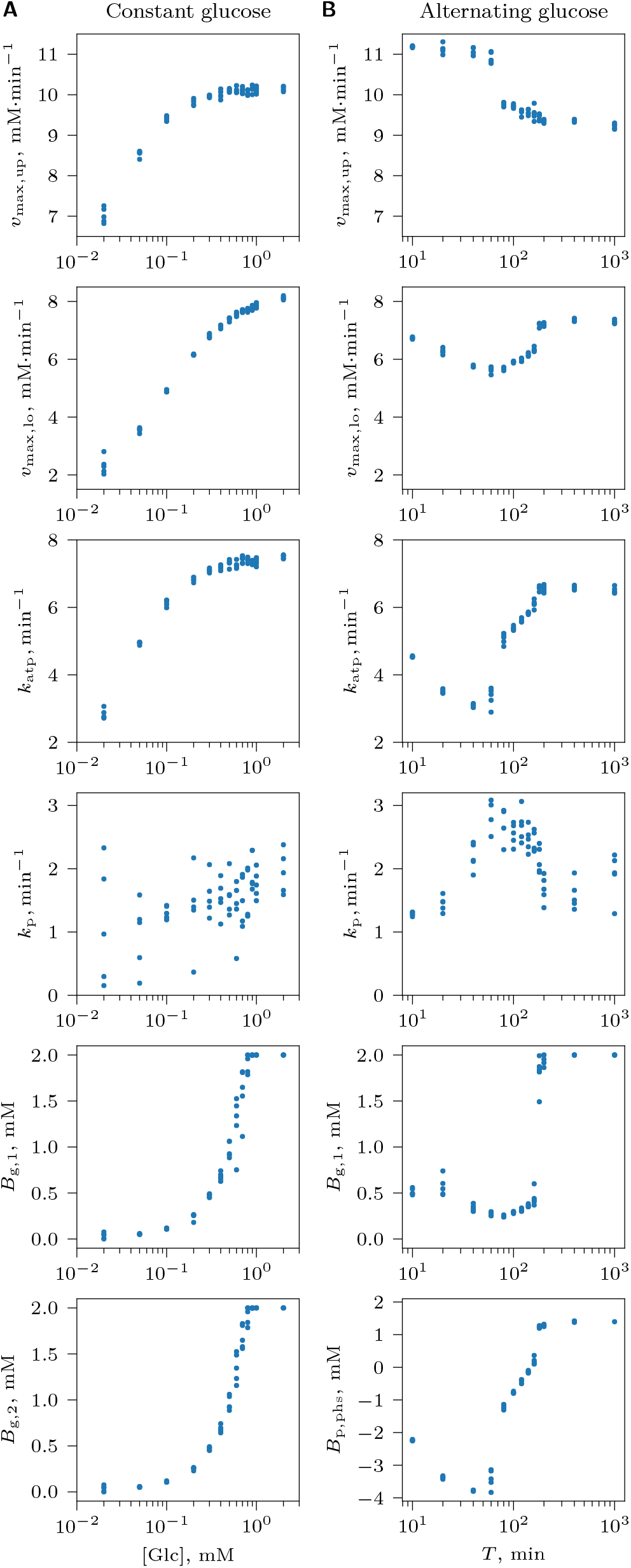
Optimization of the core glycolysis pathway in the absence of competition for glucose under NCG conditions with (A) a constant glucose supply concentration [Glc]_0_, or (B) an alternating glucose availability consisting of ON ([Glc]_0_ = 2 mM) and OFF ([Glc]_0_ = 0.01 mM) phases of equal duration with period *T* = *T*_on_+*T*_off_. Each dot represents the average of an evolving genotype parameter (*v*_max,up_, *v*_max,lo_, *k*_atp_ and *k*_p_) or a measure of balancedness at the end of an evolutionary simulation (*t*_e_). Genotype parameter averages were computed over the entire population of cells; balancedness values *B*_g,1_*, B*_g,2_ were averaged over a randomly selected subpopulation of cells that were tracked individually, and *B*_p,phs_ was calculated for the subset of tracked cells that survived through at least one ON and one OFF phase. Results of 5 replicate simulations are shown for each of the studied [Glc] and *T* value.

When comparing optimal expression patterns between environments, we observe that pathways evolve to increase the difference *v*_max,up_ − *v*_max,lo_ as glucose concentration decreases (Figure 2A). Because the risk for a cell to become imbalanced is low at low glucose concentration, and the costs of UG and LG are comparable (*w*_up_ = *w*_lo_ = 1), cells evolve higher *v*_max,up_ at the expense of *v*_max,lo_ to increase glucose uptake and thus gain a competitive advantage. As a consequence, these cells become more vulnerable to imbalanced dynamics at high glucose concentration (Figure 2A, *B*_g,1_ and *B*_g,2_). Throughout, we observe low values of *k*_p_ and a relatively high level of variation in this parameter, indicating that *k*_p_ is under weak selection. Since cells at constant glucose must show a balanced phenotype to survive, and phosphate transport is of little importance for balanced cells, *k*_p_ likely evolves to low values solely in response to weak selection for a reduction in the cost of enzyme expression.

Results are qualitatively similar when the cost of LG is much larger than that of UG (*w*_up_ = 0.1, *w*_lo_ = 1). In this case, high expression of UG enzymes can evolve to enhance glucose uptake at low glucose without major costs to the cell, while the expression of LG enzymes cannot be increased to the same level without the cell incurring prohibitive costs. By contrast, when the cost of LG is much lower than that of UG (*w*_up_ = 1, *w*_lo_ = 0.1), LG evolves high expression levels matched with the rate of UG, so that the evolved cells are balanced under any glucose concentration (Figure S2).

### Imbalanced dynamics shows fitness advantage over balanced dynamics at quickly varying glucose

Next, we studied the adaptation of the pathway to a fluctuating NCG environment with alternating [Glc]_0_ = 2 mM ON and [Glc]_0_ = 0.01 mM OFF phases of equal duration. Interestingly, cells of various balancedness coexisted in the population to form a continuum of strategies of similar fitness, from strongly balanced to strongly imbalanced (Figure 3 and Figure 4 at time *t*_me_, and Videos S1-S3). The two extremes of this balancedness continuum illustrates two radically different strategies of survival under varying glucose: upon sudden glucose availability during the ON phase, balanced cells (BCs) immediately start maintaining high ATP levels and grow, and continue to do so until the ON phase is over (Figure 3A), whereas imbalanced, “greedy” cells (ICs) do not immediately elevate ATP level, but channel all produced ATP to accumulate FBP as intracellular storage that is used up during the OFF phase to maintain a high level of ATP needed for growth (Figure 3C). The polymorphism in the population was only observed during the mutation-on segment of the simulation (i.e. between times *t*_ms_ and *t*_me_), indicating that it was caused by mutation-selection balance, a dynamic steady-state in which inferior mutants are created at the same rate as they are purged from the population by selection^20^. After mutations were stopped, only one strategy with the highest fitness survived at the final time point (Figure 4, time *t*_e_).

**Figure 3.**
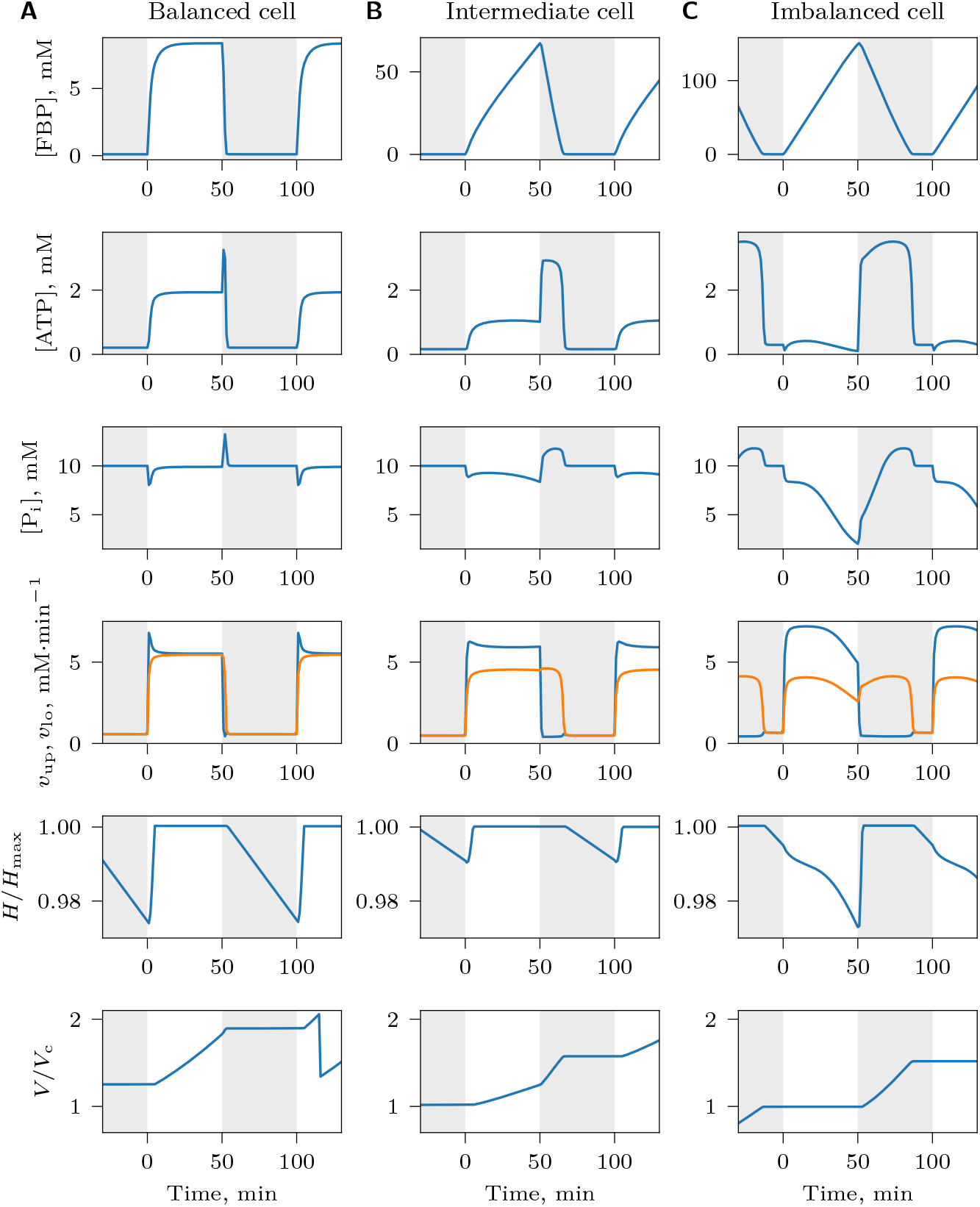
Metabolite and growth dynamics of individual cells under NCG conditions with an alternating high and low supply of glucose. Dynamics of (A) a balanced cell, *B*_p,phs_ = 1.45 mM, (B) a cell that is neither strongly balanced nor strongly imbalanced, *B*_p,phs_ = −0.03 mM, and (C) an imbalanced cell, *B*_p,phs_ = −2.13 mM. Regions with white and gray background indicate alternating glucose supply ON phase ([Glc]_0_ = 2 mM) and OFF phase ([Glc]_0_ = 0.01 mM), respectively. In the *v*_up_, *v*_lo_ plot, *v*_up_ is shown in blue and *v*_lo_ in orange. Time is shown relative to the beginning of a ON-OFF cycle. Note the different scales of FBP dynamics in (A), (B) and (C). In (A), a sudden drop in *V*/*V*_c_ indicates a cell division.

**Figure 4.**
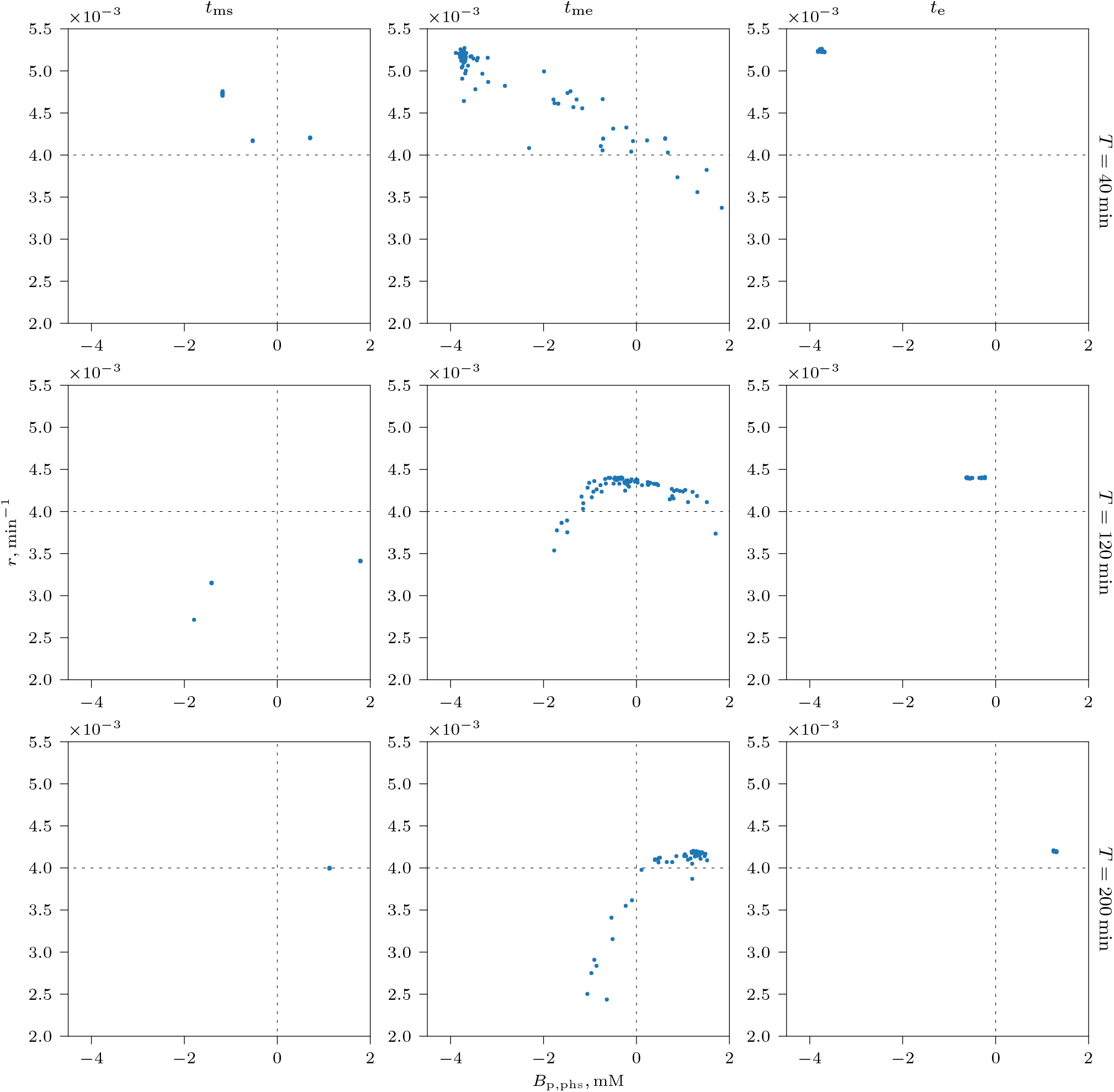
Average reproduction rate *r* of tracked cells during an environmental cycle (ON and OFF phase), plotted against their phenotypic balancedness *B*_p,phs_ at different times (*t*_ms_, *t*_me_ and *t*_e_, columns) in the NCG scenario with alternating glucose supply for different glucose pulse period *T* (rows). Points right of vertical dashed line (*B*_p,phs_ > 0) indicate balanced cells, whereas points left of the line (*B*_p,phs_ < 0) indicate imbalanced cells. The mutation-on segment of the simulation starts at time *t*_ms_ with surviving genotypes sampled from random standing genetic variation introduced at the beginning of the simulation. Mutations with small phenotypic effects then allow for a gradual evolution of the reaction rates between times *t*_ms_ and *t*_me_. Mutation is switched off again during the final segment of the simulation (between *t*_me_ and *t*_e_) so as to allow suboptimal genotypes to be purged from the population. Points with higher *r* values reflect higher cell growth rates; cells with the highest *r* values at time *t*_ms_ are the ones to survive at the end of the simulation (time *t*_e_). Therefore, *r* appears to be a good proxy for cell fitness (i.e., reproduction rate *r* minus death/removal rate), indicating that cells of different strategies do not differ markedly in removal and death rate. Videos S1-S3 show the dynamics of these plots during the whole length of simulation.

The optimal strategy depended on the glucose pulse period *T*: ICs survived at quickly varying glucose (Figure 4, *T* = 40 min and Video S1), whereas BCs were favored when the period of the environmental fluctuations was long (Figure 4, *T* = 200 min and Video S3). Periods of intermediate lengths resulted in strategies that were neither strongly balanced nor imbalanced (Figure 4, *T* = 120 min, Figure 3B and Video S2). This dependency is reflected in the evolved genotypes: the difference *v*_max,up_ − *v*_max,lo_ is large for short *T*, and small for long *T*, a feature of imbalanced and balanced genotypes respectively (Figure 2B). Evolved *k*_p_ values have smaller variation than at constant glucose and are largest at *T* values with the most strongly imbalanced cells (Figure 2B). This is consistent with a selective pressure to keep phosphate transport at an optimal level, because it plays a crucial role for FBP accumulation in ICs.

Why does the optimal strategy in a periodically fluctuating environment depend on the cycle length *T*? With fluctuating glucose availability, ICs would be expected to have higher fitness than BCs irrespective of *T*, because they could sustain larger *v*_max,up_ at the expense of *v*_max,lo_, and thus would be able to take up glucose faster than BCs. Consistent with this idea, ICs that evolved at small *T* indeed take up glucose faster than BCs that evolved at long *T*, and therefore have higher *v*_atp_ flux (Figure S4). One obvious candidate mechanism that may explain the success of BCs at large *T* is phosphate depletion: when the glucose pulse period is long, FBP accumulates to high concentrations in the cytosol, depleting phosphate reserves from the vacuole. As a result, subsequent accumulation of FBP during the ON phase becomes less efficient, limiting the potential of growth of ICs. FBP accumulation does indeed slow down with increasing FBP in the cytosol (see e.g. Figure 7C), but further analysis shows that this effect is not solely responsible for the fact that BCs outcompete ICs in slowly fluctuating environments. In particular, if we allow the network to evolve without phosphate depletion in the vacuole (*K*_vac_ → ∞), we still observe the evolution of BCs in slowly varying environments. The effect also remains if we reduce the cost of phosphate transport by an order of magnitude (*w*_p_ = 0.1), if cell health does not deteriorate (*t*_d_ → ∞) or if a different expression cost function is used (Equation 13). Instead, the observed robust advantage of BCs at large *T* derives from a difference in the timing of cell growth between BCs and ICs, allowing BCs to process more glucose during the cycle even with smaller glucose flux through the UG reaction (amount of glucose per time per volume). This asymmetry arises because the total amount of glucose converted by a cell, and therefore its fitness, is proportional to cell volume; larger cells can process more glucose per unit time. Since BCs take up glucose during the ON phase while simultaneously increasing in volume, the amount of glucose a cell can convert per unit time also increases as the ON phase progresses. Conversely, ICs take up glucose during the ON phase at constant volume; as a result, the amount of glucose a cell converts per unit time remains constant with increasing length of the ON phase. As a result, BCs ultimately do better when the period of the environmental fluctuations is large (Supplementary Text). This effect can be seen in Figure 4, where at *T* = 40 min, growth rate of ICs is higher than that of BCs, but the situation is opposite at *T* = 200 min.

Interestingly, cell balancedness also slightly increases in rapidly fluctuating environment (very small *T*, Figure 2B). This suggests that another selective pressure on ICs plays a role: although increase in *v*_max,up_ at the expense of *v*_max,lo_ will enhance glucose uptake rate and give a competitive advantage to cells, *v*_max,lo_ still has to be fast enough for all FBP accumulated during the ON phase to be fully used up during the OFF phase. This challenge becomes more difficult as *T* becomes smaller. Indeed, the *T* value below which cell balancedness slightly increases (≈ 60 min) corresponds to the point where the FBP usage time during the OFF phase approaches the length of the OFF phase (Figure S4).

One potential complication in interpreting our results is possible phenotype switching in cells with similar genotypes. In the absence of mutation, cell division does not perturb metabolite equilibria in daughter cells (Equation 7). However, a mutation upon cell division can trigger the switch of metabolic balancedness to a different state in the daughter cell, even though its genotypic balancedness is similar to that of the parent cell. To check for this problem, we compared the phenotypic and genotypic balancedness in our simulations and found that *B*_g,1_ and *B*_p,phs_ are well correlated (Figure 2B). This indicates that phenotype is largely determined by the genotype in the range of studied *T* values and that phenotypic variation of cells with similar genotypes is minimal.

### Competition for glucose can give rise to stable coexistence of balanced and imbalanced cells

After characterizing the optimization of the glycolytic pathway in response to different externally imposed glucose availability regimes (NCG conditions), we next considered the evolution of the reaction rates under chemostat conditions where cells compete for glucose and its availability changes in response to the evolving utilization strategy of the population. As a result of this feedback between evolutionary and ecological factors, selection may no longer lead to a single optimal genotype^21^.

The simulated input into the chemostat was a pulse train of glucose, consisting of a short *T*_on_ = 1 min ON phase of [Glc_0_] = 300 mM that resulted in a sharp increase in glucose in the chamber up to a few mM, followed by a longer OFF phase (either of constant or variable length) with a minimal glucose supply [Glc_0_] = 0.01 mM, during which the glucose concentration in the chamber decreased due to outflow and uptake by cells. As in our earlier simulations, we observed that short and long periods of the environmental fluctuation favored ICs and BCs respectively (Figures S5 and S7). However, intermediate values of the environmental period *T* and dilution rate *D* often resulted in stable coexistence of BCs and ICs in the population (Figure 5 and Figures S6 and S8). Contrary to what was observed in the NCG simulations, the dimorphism in the chemostat conditions did not rely on a continuous generation of new mutants, i.e., was stable at the end of the mutation-off segment of the simulation, indicating that it was not supported merely by mutation-selection balance. Instead, polymorphism was maintained by negative frequency-dependent selection, whereby two strategies can stably coexist if the fitness of each is greater when rare, a phenomenon known as protected polymorphism^20^.

**Figure 5.**
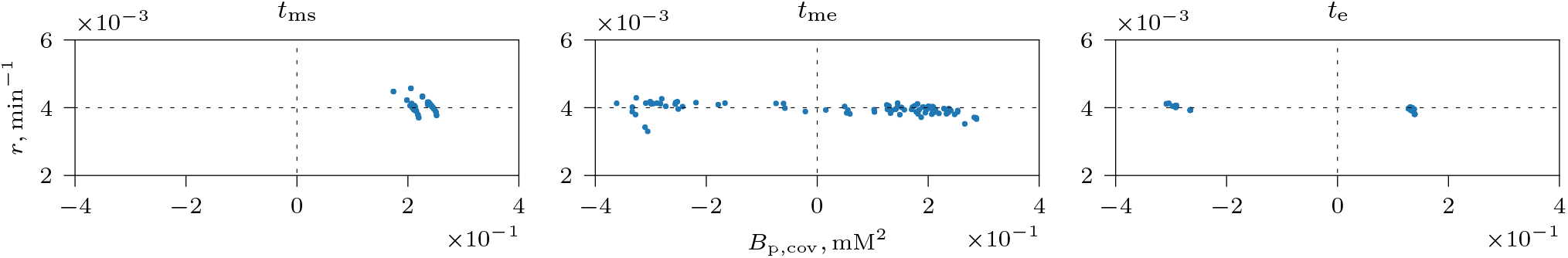
Evolution of a dimorphic population. Each dot represents the reproduction rate *r* of a tracked cell averaged over an environmental cycle plotted against its phenotypic balancedness *B*_p,cov_. Data are shown for three time points during a simulation in a chemostat with a variable OFF phase, 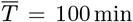, *T*_on_ = 1 min, 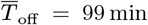, CV(*T*_off_) = 5 %, and *D* = 4 × 10^−3^ min^−1^. The initial balanced population (left, time *t*_ms_) accumulates variation created by mutation (middle, time *t*_me_). At the end of the simulation (*t*_e_), subpopulations of BCs and ICs coexist at a stable equilibrium frequency. Videos S4 shows the dynamics of this plot during the whole length of simulation.

Negative frequency dependence occurs when each strategy competes more strongly with cells of the same type than with the ones utilizing the other strategy. To demonstrate that this phenomenon is responsible for the observed coexistence, we randomly picked two genotypes from BC and IC subpopulations at the end of the simulation (shown in Figure 5, time *t*_e_, Table 2), constructed mixed populations with a range of initial fractions of BCs, *f*_b_, and ICs, 1 − *f*_b_, and then simulated their joint population dynamics in the absence of mutation. For all initial *f*_b_ values, populations restored the same equilibrium frequency of BCs (Figure 6A), except in a few cases where a high fraction of ICs (low *f*_b_) caused catastrophic dynamics and ICs were wiped out, seen as a sudden jump in *f*_b_ value to 1 (see Section *Evolution of increased imbalancedness…* below). Consistent with this evidence, the reproduction rates of ICs and BCs were observed to decrease as their fractions in the population increased (Figure 6B). Reproduction rate is a good proxy for cell fitness in our simulations, because cells rarely die and are removed from the chemostat only by outflow with constant removal rate per cell *D* that is independent of cell strategy (Equation 16).

**Table 2.** Genotypes used in no-mutation coexistence simulations. Also indicated are the expression costs of the genotypes.

|  | BC | IC |
| --- | --- | --- |
| $v_{\max, \text{up}}$ | 9.909 555 300 918 92 mM · min$^{-1}$ | 10.922 112 309 525 593 mM · min$^{-1}$ |
| $v_{\max, \text{lo}}$ | 6.973 883 285 185 177 mM · min$^{-1}$ | 5.651 433 873 885 383 mM · min$^{-1}$ |
| $k_{\text{atp}}$ | 6.141 047 310 170 09 min$^{-1}$ | 4.820 030 108 655 457 min$^{-1}$ |
| $k_p$ | 1.273 568 969 078 258 1 min$^{-1}$ | 2.070 991 048 312 316 6 min$^{-1}$ |
| $v_{\text{atp}, e}$ | 2.24 mM · min$^{-1}$ | 2.63 mM · min$^{-1}$ |

**Figure 6.**
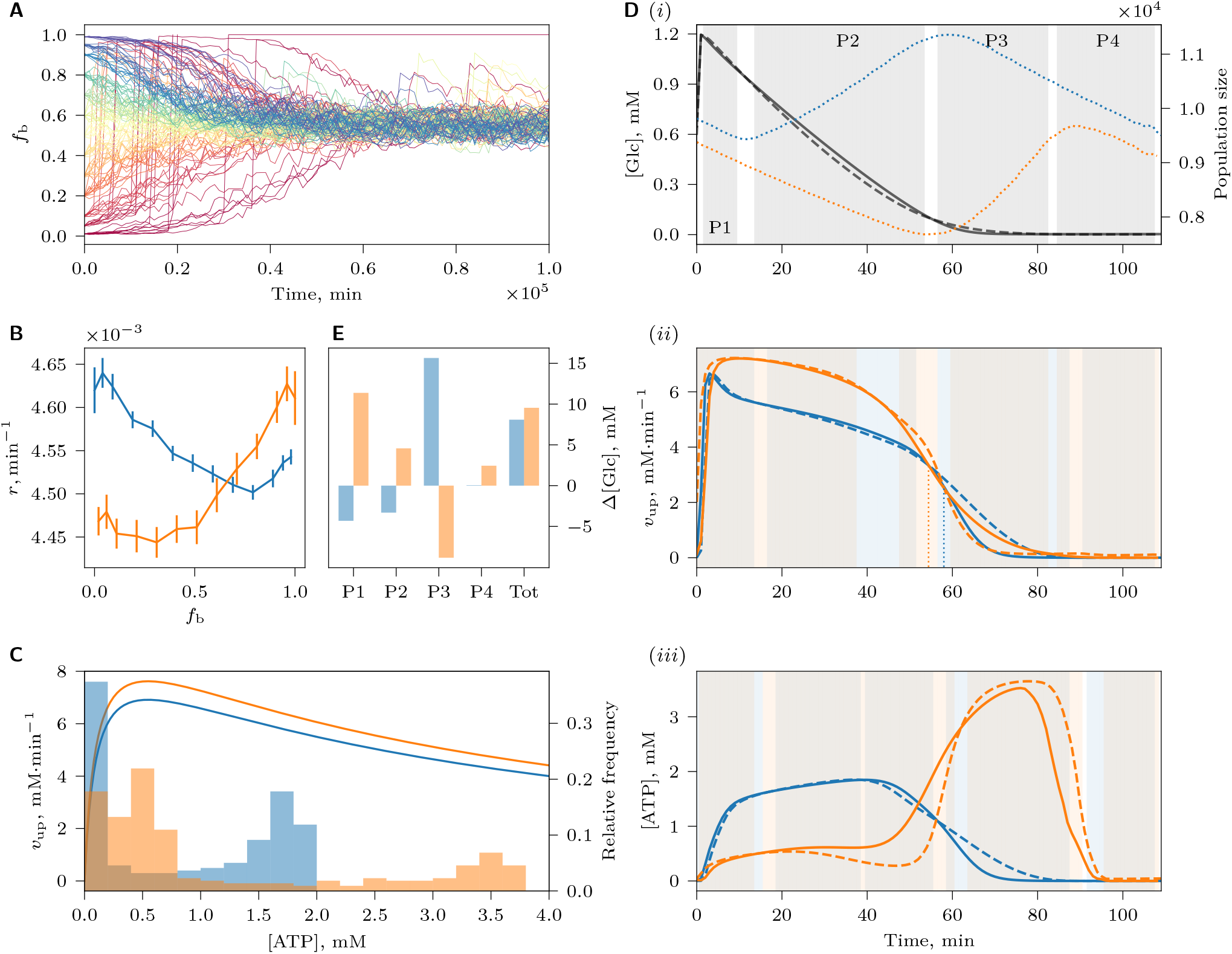
Negative frequency-dependence maintaining coexistence between BCs and ICs. (A) The joint population dynamics of two genotypes (Table 2), picked from the BC and IC subpopulations of the dimorphic population shown in Figure 5, was simulated under chemostat conditions as in Figure 5, but without mutation. Trajectories show the fraction of BCs (*f*_b_) in the population over time, for multiple different initial values of *f*_b_, indicated by color. In B, C, D(ii), D(iii) and E, data for BCs and ICs are shown in blue and orange, respectively. (B) Re-production rate of BCs and ICs as a function of their fractions in the population. Error bars indicate confidence intervals (*α* = 0.05) of the mean of reproduction rate estimates across simulation replicas. (C) Rate of UG, *v*_up_ (lines, left axis), as a function of intracellular ATP concentration (Equation 1) for BCs and ICs at [Glc] = 2 mM. Histograms (right axis) show the relative frequency distribution of intracellular [ATP] values in BCs and ICs during an environmental cycle (shown in D(iii)). (D) Average glucose, *v*_up_ and ATP dynamics during an environmental cycle (horizontal axis measures the time since the start of the glucose pulse) for a cell in a BCP (*f*_b_ = 0.99) or ICP (*f*_b_ = 0.01). (i) Average glucose concentration profile (left axis) during an environmental cycle in a BCP (solid line) and an ICP (dashed line). Gray background indicates phases P1-P4 during which the glucose concentration in BCP and ICP differ significantly (*α* = 0.05, Bonferroni adjusted). Blue and orange dotted lines indicate average population sizes of BCP and ICP respectively (right axis). Due to the difference in the timing of reproduction of BCs and ICs, BCP and ICP show different size dynamics: BCP increases in size during the first half of the cycle, when BCs reproduce, and decreases in size when reproduction stops and cells are only removed from the chemostat by the outflow. Conversely, ICP increases in size in the second half of the cycle, when ICs reproduce. (ii) Average *v*_up_ and (iii) ATP dynamics during an environmental cycle, when the focal cell type is the dominant type (solid line) or the rare type (dashed line) in the population. Because the expression cost of an IC is higher than that of a BC (Table 2), an IC has to take up more glucose than a BC to maintain the same growth rate (i.e. the area under *v*_up_ curve of an IC is larger than that of a BC). Blue (resp., orange) background indicates phases where the averages shown by blue (resp., orange) curves differ significantly; gray background highlights phases where both differ within each pair. Dotted lines in (ii) indicate time points when *v*_up_ of BCs and ICs in the population (when they one of them is dominant or rare) becomes equal as glucose is taken up. (E) Advantage of rarity measured as the differential glucose uptake per unit cell volume, Δ[Glc], compared between populations where the focal cell type is rare versus dominant. Data are shown integrated over the entire environmental cycle (Tot), as well as separately for the phases P1-P4 (here, these include both shaded areas in D(i) and half of the adjacent white space between them). Due to demographic stochasticity, estimates in (B) and (D) were obtained by averaging over many environmental cycles after [Glc] in the chemostat had reached equilibrium and where *f*_b_ has not deviated markedly from the considered value.

Frequency dependence in the chemostat arises because a population dominated by BCs (BCP) affects the profile of glucose concentration dynamics in the chemostat chamber in a different way than a population dominated by ICs (ICP) (Figure 6D(i)). As a result, glucose uptake, ATP production and the growth dynamics of the two types of cells are differentially affected in the two types of populations, such that each strategy enjoys an advantage of being rare (Figure 6D(ii) and (iii)). The fitness advantage of rarity can be quantified by comparing glucose consumption over an environmental cycle by a cell of a particular type when it is rare in the population relative to when it is dominant. This difference in glucose consumption, Δ[Glc], is proportional to the differential ATP production by the strategy and thus translates directly into a difference in reproduction rate and fitness. In Figure 6E, we show Δ[Glc] over four phases of the environmental cycle that differ in the availability of glucose between BCPs and ICPs. We observe that whenever one of the two cell types profits from being rare, the other suffers a disadvantage of rarity, and thus, overall, BCs are the superior competitor during the P3 interval of the environmental cycle, whereas ICs are the superior competitor during intervals P1, P2 and P4. The positive fitness effects of rarity, however, outweigh the negative effects for both cell types when averaged over the entire environmental cycle, creating the necessary conditions for stable coexistence by negative frequency dependence. It should be noted that the glucose consumption differentials and the resulting frequency-dependent fitness effects are rather subtle for each of the two cell types, ≈ 3 %. However, fitness differentials of such magnitude can have substantial effects over evolutionary time. For example, a strategy with a competitive advantage of 3 % is expected to spread to fixation on a time scale of 4/(3 %) ≈ 133 generations. This estimate corresponds well with the time scale for convergence to equilibrium in Figure 6A: for a reproduction rate observed in the simulations (Figure 4), 133 generations/4.5 × 10^−3^ generations · min^−1^ ≈ 0.3 × 10^5^ min.

In the remaining part of this section, we provide a detailed account of the mechanisms responsible for generating negative frequency dependence, intended for interested specialist readers. Others may skip this text and proceed to the next section. At the root of the observed frequency dependence is a feature of the core glycolysis pathway whereby the rate of UG is inhibited by high concentrations of ATP, so that *v*_up_ reaches a maximum value at an intermediate ATP concentration (Equation 1, Figure 6C). As a result, cells face a trade-off between high ATP concentration, and thus high growth rate, and fast glucose uptake. Further, it should be noted that because ICs evolved a higher expression level of UG enzymes than BCs in our simulations (Table 2, also see Figure S4), the UG rate of the IC lies above that of the BC for all ATP concentrations (Figure 6C). An additional difference between the cell types reveals itself when we compare the actual ATP concentrations that occur in BCs and ICs during an environmental cycle (blue and orange histograms; Figure 6C): where BCs operate under a regime of intermediate ATP concentrations that appears to reflect a compromise between maintaining high ATP and achieving a high rate of UG, ICs can be clearly seen to switch between two different modes of operation, one maximizing *v*_up_, the other yielding a high ATP concentration. The first mode, which is characteristic of the imbalanced state, gives an extra boost to the competitive advantage of ICs at the start of the environmental cycle, when glucose is available at high concentration. However, when glucose becomes scarce at the gradual onset of starvation, ICs switch to their second mode of operation, producing high intracellular ATP concentrations from accumulated FBP. As a result, the flux through upper glycolysis shuts down abruptly in ICs, leaving most of the remaining glucose to be consumed by BCs. Since the two types of cell are more efficient in glucose uptake at different times, each type will compete more with the same than with another type, which leads to negative frequency dependence of fitness.

Figure 6D explains how this phenomenon in turn leads to the advantage of rarity for ICs and BCs during intervals P2 and P3 respectively. Let us first focus on why ICs (orange lines in D(ii) and (iii)) do relatively better in a BCP (dashed orange) than in a population dominated by their own type (ICP; solid orange) during interval P2. It should be first noted that glucose dynamics in the chemostat chamber is determined by the glucose uptake rate of the dominant cell type, as well as the size of its population. Since ICs during interval P2 are in the low ATP, efficient glucose uptake mode, BCP consumes the available glucose somewhat slower than ICP in the first half of P2 (Figure 6D(i), P2), allowing ICs to maintain the imbalanced metabolic state for a slightly longer period of time (≈ 5 min, Figure 6D(ii), dotted lines). As a result, the decrease of *v*_up_ at the onset of starvation is delayed (Figure 6D(ii)), and low ATP concentration (characteristic of metabolic imbalance) persists for a longer period of time, so that the cell can accumulate more FBP and ultimately produce more ATP during the OFF phase (Figure 6D(iii)). (Note that glucose consumption of ICP ultimately slows down in the second half of P2, because it also depends on the ICP size, which decreases during P2, as ICs do not divide and are only removed from the population by outflow (Figure 6D(i), orange dotted line); glucose concentration therefore equalizes between BCP and ICP at the end of P2). After the switch of ICs to high ATP, slow glucose uptake mode (interval P3) as glucose concentration decreases, BCs become more efficient in glucose uptake, and therefore BCP reduces the glucose concentration faster than ICP. As a result, BCs (blue lines in D(ii) and (iii)) do better when they are rare in a population dominated by ICs (dashed lines), as the remaining glucose is only very slowly consumed by ICs, but mostly is taken up by BCs. By contrast, in a BCP (solid lines), BCs compete for glucose with other cells of the same type right until little glucose is left.

The advantage of rarity of ICs during intervals P1 and P4 manifests itself through a different mechanism. At the end of the cycle (during P4), glucose concentration in the environment is very low, and ICs have already used up all their FBP. As a result, ATP concentration in both cell types is very low. Because ICs in this state take up the remaining scraps of glucose a little faster than BCs (*v*_up_ is higher for ICs close to the point [ATP] = 0, Figure 6D), ICs in BCP enjoy a slightly larger glucose concentration, and therefore a slightly larger ATP concentration, leading to the advantage of rarity. Due to the autocatalytic nature of the glycolytic pathway (i.e., the pathway needs ATP investment to generate more ATP), this will allow ICs in BCP to restart glucose uptake slightly faster upon glucose availability at the beginning of the next cycle (P1), giving them a fitness advantage.

### Evolution of increased imbalancedness renders populations vulnerable to catastrophic collapse

In the intermediate range of studied *D* and *T* values where the fittest strategy transitions from balanced to imbalanced (Figures S5 to S8), populations were often observed to exhibit catastrophic events whereby population size collapsed and then recovered (Figure 7A and B, also Figure 6A). Interestingly, during a catastrophe, the fraction of ICs in the population drops and BCs become dominant, but in between two adjacent catastrophic events the fraction of ICs increases as they are more competitive than BCs (Figure 7B and Figure S11). These eco-evolutionary cycles result from a vulnerability of ICPs: stochastic decrease in the population size can temporarily elevate the concentration of glucose in the chemostat (see *Model and methods*), forcing ICs to spend more time in the imbalanced state accumulating FBP and less time processing FBP to maintain high ATP needed for reproduction. As a result, FBP increasingly accumulates in ICs over many environmental cycles because it cannot be fully processed, and the reproduction rate of ICs decreases (Figure 7B), leading to a further decrease in the population size and elevation of the glucose level. In ICPs with strongly ICs that are particularly prone to accumulate more FBP than they can handle, this positive feedback can easily escalate into a catastrophic collapse of the population, where most of the ICs are lost. When such a catastrophe occurs, the population can survive if it still contains a small subpopulation of BCs that survived the competition with ICs. BCs profit from the increased glucose concentration by reproducing faster, causing the population size and the glucose concentration to be restored to their normal levels. The surviving population is dominated by BCs, which, however, create ideal conditions for more competitive imbalanced strategies to evolve. ICs, either the ones that survived the catastrophe, or newly generated mutants, will therefore increase in frequency and evolve to become more imbalanced (i.e. competitive) over time, causing the cycle to repeat. Catastrophic dynamics thus depends on the polymorphism in the population: ICs cause the collapse of the population, but only BCs can restore it. Vice-versa, catastrophes also appear to be a mechanism by which polymorphism is maintained, as they are crucial to prevent ICs from completely overtaking the population.

**Figure 7.**
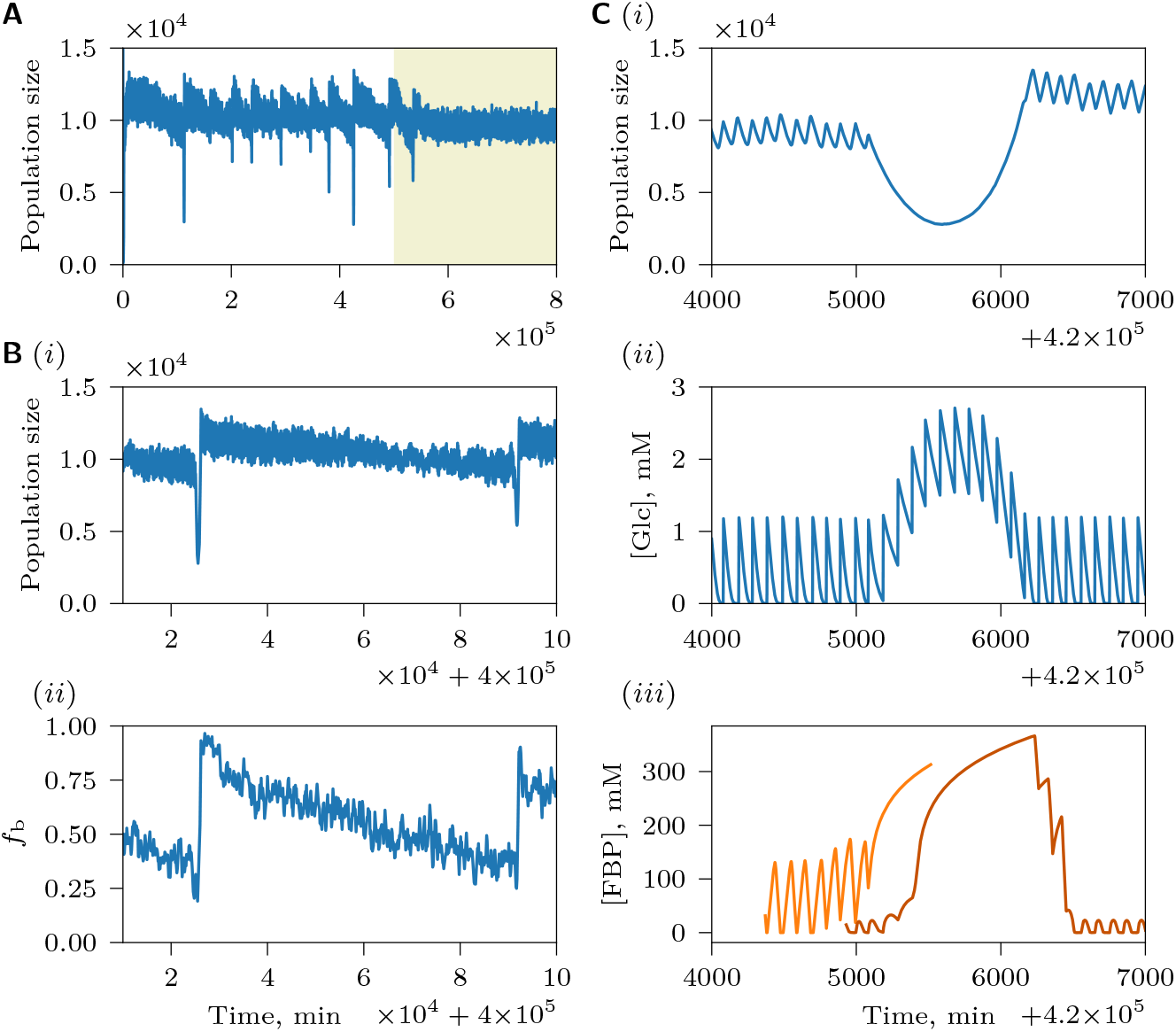
Catastrophic dynamics of the population showed in Figure 5. (A) Population size during the whole length of the simulation. Yellow background indicates the mutation-off segment of the simulation. (B) (i) Population size between two catastrophic events and (ii) corresponding fraction of BCs of tracked cells, *f*_b_, where BC is defined as having a positive *B*_p,cov_. An apparently decreasing population size between catastrophic events shown in (A) and (B)(i) is due to the difference in the timing of reproduction of BCs and ICs (Figure 6D(i)). Just after the catastrophe, the population is dominated by BCs and therefore population reaches larger sizes than immediately before the catastrophic event, when the population is dominated by ICs. (C) A close-up of a catastrophe: (i) population size, (ii) glucose dynamics and (iii) FBP dynamics of two example cells, strongly imbalanced (*B*_g,1_ = 0.15 mM, light orange) and weakly imbalanced (*B*_g,1_ = 0.55 mM, dark orange). The strongly imbalanced cell begins accumulating large amounts of FBP earlier than the weakly imbalanced cell. One cell is removed by outflow in the middle of the catastrophic event, whereas the other survives the catastrophe and recovers.

Interestingly, after mutations are stopped, populations were always observed to undergo only one catastrophe at most (Figure 7A), suggesting that *de novo* mutation might be needed to reintroduce ICs in the population after a catastrophe for continued population cycles. An additional role of mutation could be that it fuels the gradual replacement of weakly ICs by stronger imbalanced, more competitive ones, which increase the vulnerability of the population to collapse. Moreover, newly generated mutants could introduce additional randomness to the system, thus weakening the stabilizing force of negative frequency-dependence and increasing the probability of a catastrophe. Both hypotheses are supported by the observation that reducing the mutation rate decreases the frequency of catastrophes (Figure S10A and B). The role of randomness is further emphasized by the fact that the frequency of catastrophes is also reduced in the environment with consistent glucose availability (constant *T*_off_, Figure S10C). Another possibility is that continuously generated mutants affect glucose profile in the chemostat in such a way as to add a slight fitness advantage to ICs. According to this scenario, during the mutation-on segment of the simulation the fraction of ICs would tend to increase, periodically causing catastrophes, but the absence of mutants would make fitness of ICs and BCs more equal, and protected polymorphism would not allow the fraction of ICs to deviate too much from equilibrium to cause a catastrophe. This possibility is supported by the finding that the genotypes with fitness advantage during the mutation-on segment are different from the ones during the mutation-off segment (Figure S11).

## Discussion

Upon transition to high glucose, 7 % of WT yeast cells enter a non-viable state of imbalanced glycolysis, whereby UG outpaces LG and glycolytic intermediates accumulate at low ATP^3^. Computational modeling studies suggested that the two states, balanced and imbalanced, are an inherent feature of glycolysis: the pathway can be pushed towards either one of the alternative states by spontaneous heterogeneity in metabolite concentrations or enzyme levels among isogenic cells^3^. Furthermore, it has also been shown that the propensity of the simulated yeast glycolysis pathway to enter the imbalanced state can be modified by slowing down UG, speeding up LG or phosphate transport from the vacuole. In this study we address the question why yeast cells do not employ available mechanisms, such as increasing the constitutive expression of LG or phosphate transport enzymes, to minimize or entirely eliminate the risk of developing metabolic imbalance. Although it is conceivable that the cost of these mechanisms to the population do not weigh up against the substantial benefit of rescuing 7% of cells, we propose an alternative hypothesis, whereby WT yeast are prone to imbalanced glycolysis because they are evolutionarily optimized for scarce or varying glucose. Our simulations support this hypothesis: since the likelihood of entering the imbalanced state decreases with decreasing glucose concentration, model cells that evolve under scarce glucose exhibit higher expression of UG enzymes at the expense of LG enzymes to enhance glucose uptake and thus gain a competitive advantage without the risk of becoming imbalanced. However, the adaptation to scarce resource makes them more vulnerable to imbalanced dynamics when glucose is available abundantly.

Furthermore, in variable environments with rapidly fluctuating glucose levels, the seemingly maladaptive imbalanced state provided a clear fitness advantage over balanced metabolism: during the period of glucose abundance, ICs quickly accumulated FBP as intracellular storage that was then consumed during the period of scarcity to maintain high ATP and reproduce. Note that this benefit can only materialize if the imbalanced state is reversible: experimental observations of yeast cells trapped in the imbalanced state show that they are viable for around 7 h and can resume growth on galactose when glucose is removed^3^. Our model indicates that imbalanced metabolism can be favored by selection because ICs can evolve higher levels of UG enzymes at the expense of LG enzymes to secure the resource faster than BCs without compromising the overall performance of the glycolysis pathway. However, in environments with long periods of resource scarcity, ICs lost their fitness advantage over BCs. Since glucose uptake and cell growth are separated in time in ICs, and because the amount of glucose taken up by a cell depends on its volume, ICs do not increase their total capacity for processing glucose as they are securing resources from the environment. BCs, on the other hand, show a clear accelerating growth pattern: because BCs grow at the same time as they take up glucose, increase in cell volume immediately translates into an increase in the total metabolic capacity of the cell. As a consequence, ICs grow less efficiently than BCs when the environmental fluctuations are slow. One way to experimentally test the predictions of our model is to evolve yeast under different glucose availability regimes and determine the fraction of evolved cells that become imbalanced upon a transition to high glucose^3^. In addition, the expression levels of UG and LG enzymes in evolved strains could be quantified and compared against the patterns predicted by our model.

As mentioned above, stochastic phenotype determination, triggered by random fluctuations in metabolic state, provides a mechanism that explains the co-occurrence of balanced and imbalanced WT cells in an isogenic population upon the transition to excess glucose^3^. Our model suggests two other mechanisms that can also support a phenotypic polymorphism of BCs and ICs, which may be particularly important under natural conditions in a genetically variable population. First, our simulations show that both balanced and imbalanced dynamics represent viable strategies in an environment where the availability of glucose varies over time. Although, in any particular environment, one of the two strategies typically enjoys a competitive advantage over the other, the fitness differences between them are often small, such that substantial variation can be maintained under mutation-selection balance, whereby the rate at which less fit mutants are eliminated equals to the rate of their creation by mutation^20^. In our simulated populations, one type of cell is easily produced from another type by mutation, and since the evolvable parameters represent expression levels of enzymes, it is feasible that similar conversions could easily occur in natural populations.

The second mechanism that allows for phenotypic variation is negatively frequency-dependent selection, which can support the emergence and stable coexistence of discrete clusters of genetically differentiated BC and IC types. This protected polymorphism arises in a chemostat regime where cells compete for glucose, and their utilization strategy influences the resource concentration in the chemostat chamber. This establishes an ecological feedback: by consuming glucose in different ways, BCs and ICs induce a different dynamic of the glucose concentration, which, in turn, affects the two competing strategies in different ways. In fact, BCs create conditions favorable for the growth of ICs, and *vice versa*, such that each type enjoys the advantage of rarity, and thus diversity is maintained. Negative frequency-dependent selection has been previously experimentally demonstrated in yeast populations in multi-resource environments^22,23^. Our simulations, however, point to the possibility of protected polymorphism in a single resource chemostat environment. One way to experimentally demonstrate this could be to establish under what conditions already evolved BCs and ICs can stably coexist in a chemostat. Such experiments are lacking because laboratory studies generally work with well-characterized genetically monomorphic populations.

A further unanticipated phenomenon highlighted by our model is that a population of coexisting BCs and ICs under varying glucose can exhibit catastrophic collapses, often followed by a recovery. A prerequisite to a catastrophe is an increase in the fraction of strongly ICs in the population due to their competitiveness. However, ICs in such a population become vulnerable to falling into a self-sustaining state of accumulating more FBP than they can use to produce ATP, which reduces their efficiency of growth, causes a drop in the population size with a concomitant increase in glucose concentration in the environment that pushes even more ICs into persistent imbalance. The recovery of the population depends on the presence of BCs, either surviving ones or newly generated mutants, that benefit from the increased glucose concentration in the environment. By restoring the normal glucose level, however, BCs create ecological conditions in which ICs are competitively superior, setting the stage for the cycle to repeat itself. Therefore, the recurrent catastrophic collapse and recovery of the population requires a polymorphism of balanced and imbalanced cells, but also helps to maintain their dynamic coexistence. Although in our simulations the recurrent catastrophes require mutational pressure, it is feasible that, under some conditions, they could occur without mutational pressure and be the only mechanism to maintain polymorphism. Such a process would be akin to protected polymorphism with the difference that the decrease of fitness of dominant ICs would be delayed and dependent on chance, i.e. would only happen when a catastrophe is triggered by stochastic increase of glucose concentration due fluctuations in the population size.

The catastrophic eco-evolutionary dynamics observed in our simulations bears similarity to the phenomena of the tragedy of the commons and evolutionary suicide, particularly in cases when the population does not recover after a collapse. The tragedy of the commons occurs when individual-level adaptations driven by natural selection maximizes fitness relative to other individuals at the expense of a public good, which can result in decrease of mean population fitness (or a proxy thereof, such as overall offspring production)^24^. The resulting decrease in size can make the population vulnerable to extinction due demographic or environmental stochasticity, or, alternatively, if the disturbance pushes the population into a different stable state associated with a different regime of selection (e.g. past a bifurcation point), the population may undergo evolutionary suicide, i.e. can be driven towards extinction deterministically by natural selection^25,26^. In the context of the current model, high resilience of the population to fluctuations in glucose concentration, and thus to catastrophes, can be considered as public good for ICs. Yet, their individual-level adaptations, driven by the selective pressure to increase competitiveness by becoming more imbalanced, undermines this common good, pushing the population ever closer towards the brink of collapse.

Overall, our study demonstrates that a highly simplified metabolic network, without even considering its genetic regulation, is sufficiently flexible to encapsulate a dynamic feedback between metabolic adaptation and resource availability and that their interplay, in turn, gives rise to population level phenomena, such as the maintenance of alternative strategies or population cycles that shape selection on the metabolic network. Here, we have shown how considering this eco-evolutionary perspective sheds new light on the prevalence of substrate-accelerated death in yeast. We expect that it will do similarly well for explaining other seemingly maladaptive aspects of cellular metabolism.

## Acknowledgments

AJ and GSvD thank David Ekkers for inspiring discussions. This work was financially supported by the Netherlands Organisation for Scientific Research (NWO Vidi Grant 864.11.012 to GSvD) and the European Research Council (ERC Starting Grant 309555 to GSvD).

