## Supplementary material for "Selection for rapid uptake of scarce or fluctuating resource explains vulnerability of glycolysis to imbalance"

### Why BCs have higher fitness than ICs in slowly variable environments

Let us consider the case in NCG scenario with $t_\mathrm{d} \to \infty$, i.e. when cell health does not deteriorate and therefore the cell continuously remains at its maximum health (Figure S3C). Figure 4 suggests that the cell reproduction rate $r$, equal to the average fractional volume increase rate during the environmental ON-OFF cycle, is a good proxy for cell fitness:

$$r = \frac{1}{T}\int_0^T \frac{V'}{V}\,dt\,, \tag{21}$$

where $T_\mathrm{on} = T_\mathrm{off} = \frac{1}{2}T$, and the cycle is assumed to start at time $t = 0$. Since our aim is the comparison of fitness between two different strategies (i.e., the sign of difference), we will employ a simpler measure, the average total volume increase rate during the environmental cycle,

$$W = \frac{V(T) - V(0)}{T} = \frac{1}{T}\int_0^T V'\,dt\,, \tag{22}$$

because it is monotonically related to the reproduction rate and fitness proper due to both cell types having the same standard volume $V_\mathrm{c}$. If we define

$$v^+_{\mathrm{atp,g}} = \begin{cases} v_{\mathrm{atp,g}} & \text{if } v_{\mathrm{atp,g}} \geq 0\,,\\ 0 & \text{if } v_{\mathrm{atp,g}} < 0\,,\end{cases} \tag{23}$$

then, from Equation 10,

$$V' = u_\mathrm{g} v^+_{\mathrm{atp,g}} V\,, \tag{24}$$

$$W = \frac{u_\mathrm{g}}{T}\int_0^T v^+_{\mathrm{atp,g}} V\,dt\,. \tag{25}$$

$W$ is thus proportional to the average rate of ATP usage for growth per cell during the environmental cycle, and therefore increases with the increasing average rate of glucose uptake per cycle.

**Balanced cells.** A BC grows during the ON phase, when [ATP], $v_\mathrm{atp}$ and [FBP] are approximately constant, maintained by the influx of glucose into the cell (Figure 3A). During the OFF phase, cell volume does not increase, and therefore

$$W_\mathrm{b} = \frac{u_\mathrm{g}}{T}\int_0^{\frac{1}{2}T} v^+_{\mathrm{atp,g}} V\,dt = \frac{u_\mathrm{g}}{T} v^+_{\mathrm{atp,g}} \int_0^{\frac{1}{2}T} V\,dt\,. \tag{26}$$

From Equation 24 it follows that a cell grows exponentially when the flux $v^+_{\mathrm{atp,g}}$ is constant:

$$V = V_0 e^{u_\mathrm{g} v^+_{\mathrm{atp,g}} t}\,, \tag{27}$$

where $V_0$ is the initial cell volume at time $t = 0$, and

$$\int_0^{\frac{1}{2}T} V\,dt = V_0 \int_0^{\frac{1}{2}T} e^{u_\mathrm{g} v^+_{\mathrm{atp,g}} t}\,dt = \frac{V_0}{u_\mathrm{g} v^+_{\mathrm{atp,g}}} e^{u_\mathrm{g} v^+_{\mathrm{atp,g}} t}\Big|_0^{\frac{1}{2}T} = \frac{V_0}{u_\mathrm{g} v^+_{\mathrm{atp,g}}}(e^{\frac{1}{2}u_\mathrm{g} v^+_{\mathrm{atp,g}} T} - 1)\,. \tag{28}$$

From Equations 24 and 26,

$$W_\mathrm{b} = \frac{V_0}{T}\left(e^{\frac{1}{2}u_\mathrm{g} v^+_{\mathrm{atp,g}} T} - 1\right)\,. \tag{29}$$

$W_\mathrm{b}$ will thus increase exponentially with increasing $T$, until $T$ becomes so large that a cell division occurs. This happens because a BC takes up glucose as it grows; since a larger cell can take up more glucose, the cell speeds up its absolute glucose consumption with increasing size, and therefore the average glucose uptake rate per cell will increase with increasing $T$.

**Imbalanced cells.** Our simulations show that an IC does not grow during the ON phase, but accumulates FBP that is used up for cell growth during the OFF phase (Figure 3C). Because in a competitive IC FBP is completely used up before the end of the OFF phase, $v^+_{\mathrm{atp,g}}$ is not constant throughout the phase but will drop before the phase is over. Equation 25 can be evaluated by assuming that during the OFF phase, every FBP molecule converted by LG will produce 4 ATP molecules that will be used up by the ATPase reaction (Figure 1). As FBP concentration during the conversion is large, LG is saturated, $v^+_{\mathrm{atp,g}} = \mathrm{const}$, and a constant fraction of generated ATP will be used for cell maintenance costs $v_{\mathrm{atp,c}}$. If $q = \frac{v^+_{\mathrm{atp,g}}}{v_{\mathrm{atp}}}$ represents the fraction of ATP used for cell growth, then

$$W_{\mathrm{i}} = \frac{u_{\mathrm{g}}}{T} \int_{\frac{1}{2}T}^{T} v^+_{\mathrm{atp,g}} V \, dt = \frac{u_{\mathrm{g}} q}{T} \int_{\frac{1}{2}T}^{T} v_{\mathrm{atp}} V \, dt = \frac{4 u_{\mathrm{g}} q}{T} \int_{\frac{1}{2}T}^{T} v_{\mathrm{lo}} V \, dt \,, \tag{30}$$

i.e., $W_{\mathrm{i}}$ is proportional to $\frac{4}{T} \int_{\frac{1}{2}T}^{T} v_{\mathrm{lo}} V \, dt$, the rate of ATP generation by LG per cell during the OFF phase. Because the amount of FBP in the cell, $cV$, where $c$ is the concentration of FBP, decreases only due to LG, $(cV)' = c'V$, and

$$\int_{\frac{1}{2}T}^{T} v_{\mathrm{lo}} V \, dt = \int_{\frac{1}{2}T}^{T} (-c') V \, dt = -\int_{\frac{1}{2}T}^{T} (cV)' \, dt = c(\frac{1}{2}T)V(\frac{1}{2}T) - c(T)V(T) \,. \tag{31}$$

Further, (i) FBP concentration at the beginning of the OFF phase is $c(\frac{1}{2}T) = \frac{1}{2}aT$, where $a$ is FBP accumulation rate during the ON phase, (ii) all FBP is used up during the off phase, i.e. $c(T) = 0$, and (iii) $V(\frac{1}{2}T) = V_0$, because the cell does not grow during the ON phase. As a result,

$$\int_{\frac{1}{2}T}^{T} v_{\mathrm{lo}} V \, dt = \frac{1}{2} a T V_0 \,, \tag{32}$$

and, from Equations 30 and 32,

$$W_{\mathrm{i}} = 2 u_{\mathrm{g}} q a V_0 \,, \tag{33}$$

i.e. $W_{\mathrm{i}}$ is independent of $T$. This happens because ICs take up glucose during the ON phase at constant volume; the increase in $T$ does not increase the average glucose uptake rate per cell.

This analysis illustrates why BCs have higher fitness than ICs at large $T$: as $T$ increases, average glucose uptake rate of ICs during the cycle remains the same, whereas that of BCs increases. Plotting $W_{\mathrm{b}}$ and $W_{\mathrm{i}}$ for representative parameter values observed in our simulations shows that $W_{\mathrm{i}}$ becomes smaller than $W_{\mathrm{b}}$ at $T$ values of several hundreds of minutes, in good agreement with the simulation results (Figure S1). The difference in these predicted and the observed $T$ values can be attributed to cell health dynamics and cell divisions that are not accounted for in this analysis.

We can also find the reproduction rate (the proxy of fitness in our study) of a BC directly:

$$r_{\mathrm{b}} = \frac{1}{T} \int_0^T \frac{V'}{V} \, dt = \frac{u_{\mathrm{g}}}{T} \int_0^{\frac{1}{2}T} v^+_{\mathrm{atp,g}} \, dt = \frac{u_{\mathrm{g}}}{T} v^+_{\mathrm{atp,g}} \cdot \frac{1}{2} T = \frac{1}{2} u_{\mathrm{g}} v^+_{\mathrm{atp,g}} \,. \tag{34}$$

This shows that the fitness of a BC is constant, independent of $T$. Since we have established that the fitness of ICs becomes smaller than that of BCs at long $T$, it can be concluded that the fitness of ICs decreases with increasing $T$. In other words, while a BC increases its total metabolic capacity per cell, and therefore the rate of average absolute volume increase $W_{\mathrm{b}}$ as the cell grows, so that the rate of fractional volume increase remains the same, the metabolic capacity of an IC, and thus $W_{\mathrm{i}}$ remains the same as the cell grows, resulting in the decreasing rate of fractional volume growth.

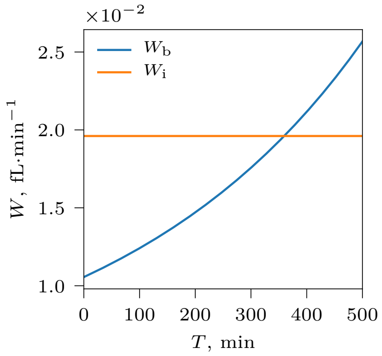


**Figure S1.** Average volume increase rate of a BC during an environmental cycle, $W_b$ (blue, Equation 26), becomes larger than that of an IC, $W_i$ (orange, Equation 30) as the period of environmental fluctuations $T$ increases. The plots are shown for representative parameter values observed in NCG simulations: $v^+_{atp,g} = 6.3\,\text{mM} \cdot \text{min}^{-1}$, $q = 0.77$, $a = 3.8\,\text{mM} \cdot \text{min}^{-1}$, $V_0 = V_c$.

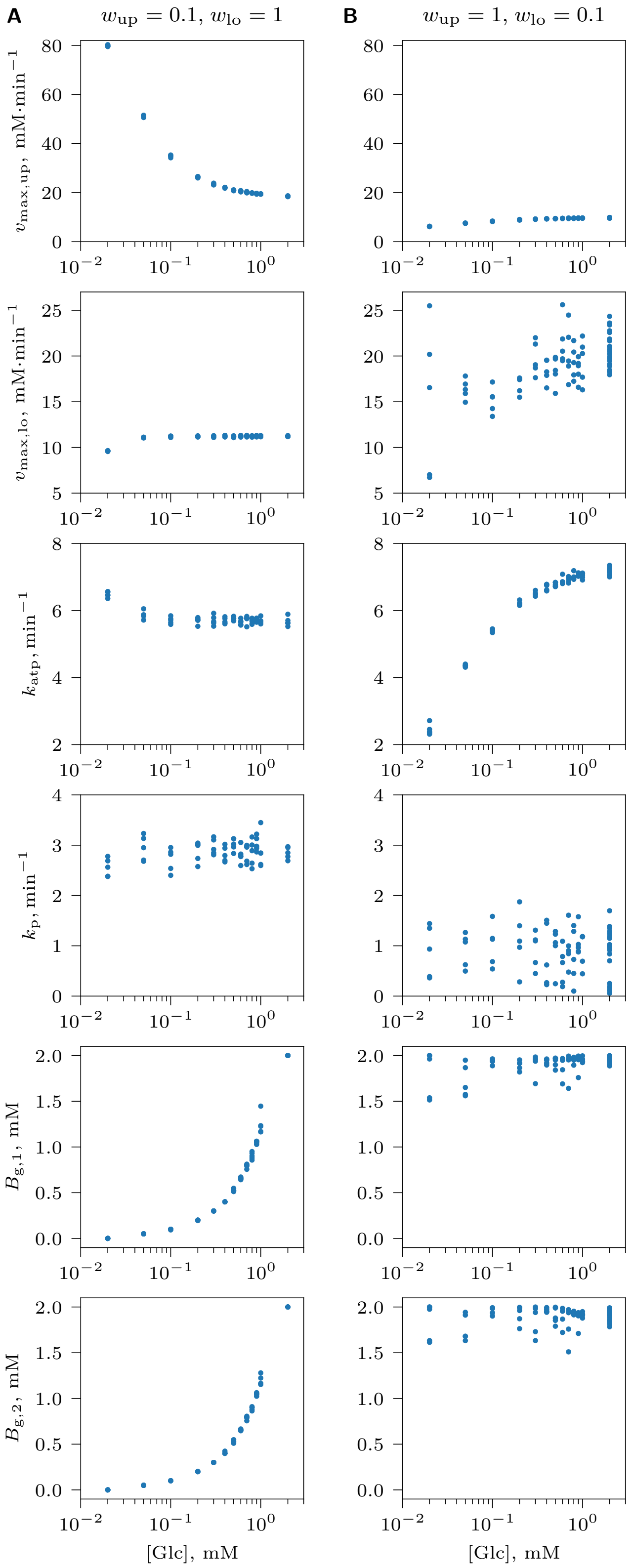


**Figure S2.** Optimization of the core glycolysis pathway in the absence of competition for glucose under NCG conditions with a constant glucose supply concentration, when the costs of UG and LG differ markedly. (A) UG is less costly, $w_{\text{up}} = 0.1$, $w_{\text{lo}} = 1$, (B) LG is less costly, $w_{\text{up}} = 1$, $w_{\text{lo}} = 0.1$. Each dot represents the average of an evolving genotype parameter ($v_{\text{max,up}}$, $v_{\text{max,lo}}$, $k_{\text{atp}}$ and $k_{\text{p}}$) or a measure of balancedness at the end of an evolutionary simulation ($t_{\text{e}}$). Genotype parameter averages were computed over the entire population of cells; balancedness values $B_{\text{g},1}, B_{\text{g},2}$ were averaged over a randomly selected subpopulation of cells that were tracked individually. Results of 5 replicate simulations are shown for each of the studied [Glc] value.

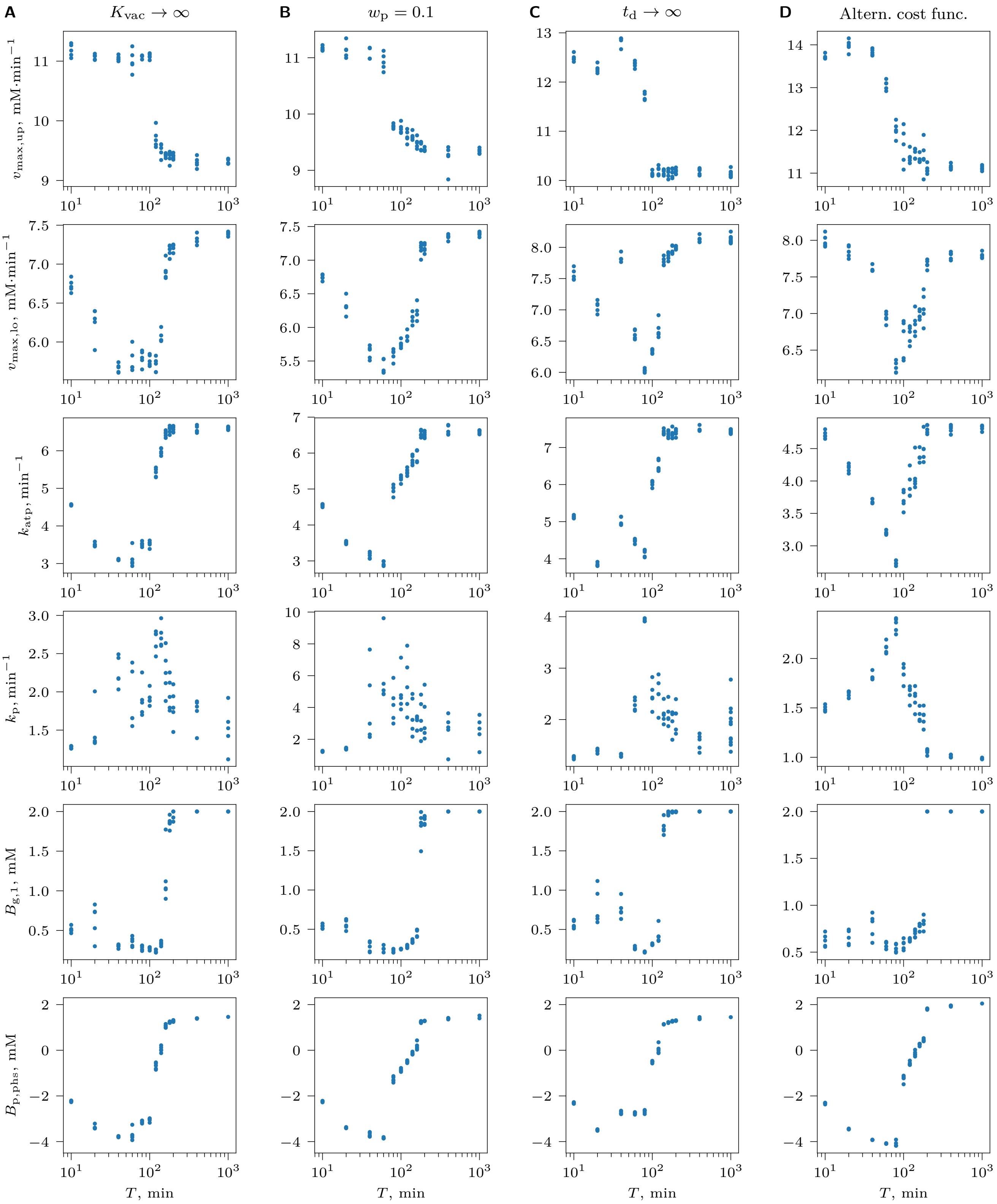


**Figure S3.** Optimization of the core glycolysis pathway in the absence of competition for glucose under NCG conditions with an alternating glucose availability consisting of ON ($[Glc]_0 = 2$ mM) and OFF ($[Glc]_0 = 0.01$ mM) phases of equal duration with period $T = T_{on} + T_{off}$, when (A) no depletion of phosphate from the vacuole occurs, $K_{vac} \to \infty$, (B) the cost of phosphate transport is small, $w_p = 0.1$, (C) cell health does not decrease, $t_d \to \infty$, and therefore cells do not die, and (D) the alternative cost function is used (Equation 13). Each dot represents the average of an evolving genotype parameter ($v_{max,up}$, $v_{max,lo}$, $k_{atp}$ and $k_p$) or a measure of balancedness at the end of an evolutionary simulation ($t_e$). Genotype parameter averages were computed over the entire population of cells; balancedness values $B_{g,1}, B_{g,2}$ were averaged over a randomly selected subpopulation of cells that were tracked individually, and $B_{p,phs}$ was calculated for the subset of tracked cells that survived through at least one ON and one OFF phase. Results of 5 replicate simulations are shown for each of the studied [Glc] and $T$ value.

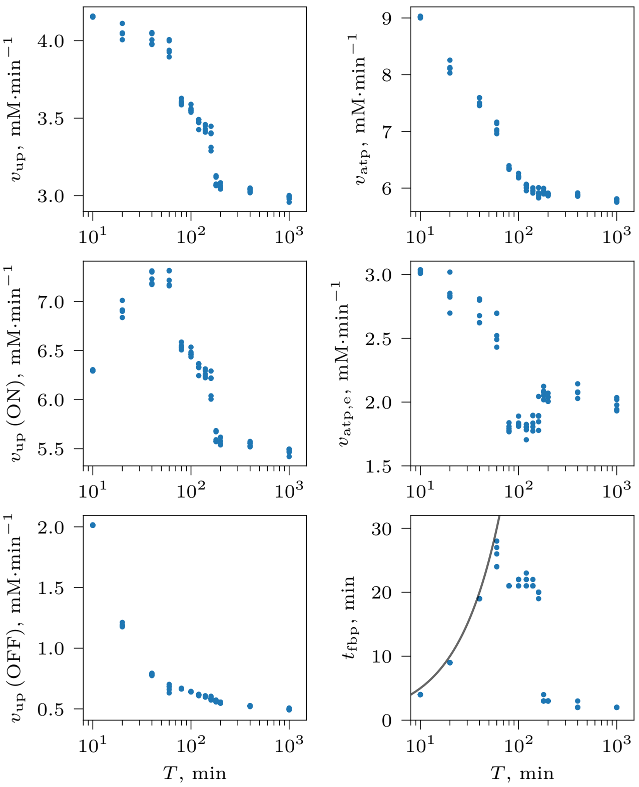


**Figure S4.** Optimization of the core glycolysis pathway in the absence of competition for glucose under NCG conditions with an alternating glucose availability consisting of ON ($[\mathrm{Glc}]_0 = 2\,\mathrm{mM}$) and OFF ($[\mathrm{Glc}]_0 = 0.01\,\mathrm{mM}$) phases of equal duration with period $T = T_{\rm on} + T_{\rm off}$ (see Figure 2B). Each dot represents the average of a flux $v_{\rm up}$, $v_{\rm atp}$, $v_{\rm atp,e}$ over an environmental cycle, or the time when [FBP] falls below 5 mM (i.e., is used up) after the beginning of the OFF phase, $t_{\rm fbp}$. The averages were computed over a randomly selected subpopulation of cells that were tracked individually at the end of an evolutionary simulation ($t_{\rm e}$) and that survived through at least one ON and one OFF phase. Results of 5 replicate simulations are shown for each of the studied $T$ value. The black line indicates $t_{\rm fbp} = T_{\rm off} = \frac{1}{2}T$, i.e. where FBP is used up exactly at the end of the OFF phase.

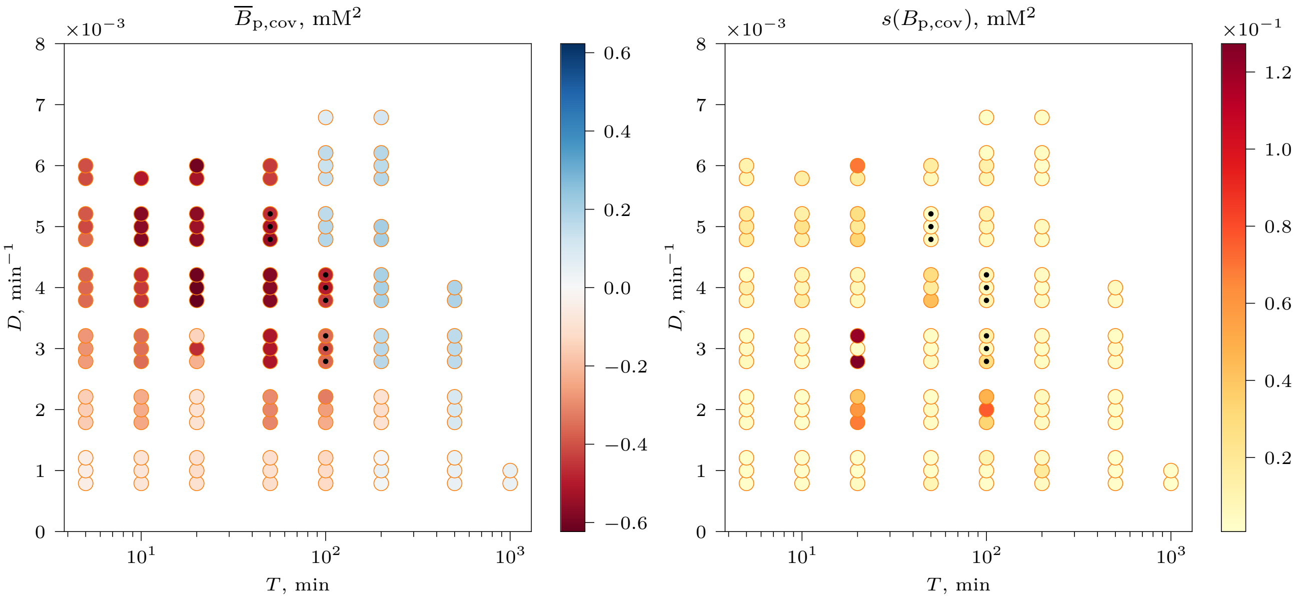


**Figure S5.** Overview of the average phenotypic balancedness $\overline{B}_{\mathrm{p,cov}}$ and its standard deviation $s(B_{\mathrm{p,cov}})$ of evolved tracked cells (colorbar) at the end of an evolutionary simulation ($t_{\mathrm{e}}$) with constant $T_{\mathrm{off}}$ for a wide range of glucose pulse period $T$ and chemostat flow rate $D$ values. A group of three or less circles represents three replicate simulations for the same pair of $T$ and $D$ values (i.e., circles have been displaced in the vertical direction for the purpose of visualization; a $D$ value of the group is at the closest tick mark on the vertical axis). A circle in the plot is absent when the population did not survive to the simulation end, or the population size $N(t_{\mathrm{e}}) < 1000$. Black dots indicate simulations where catastrophic dynamics was observed (see Section *Evolution of increased imbalancedness...*). High variation in balancedness of cells at $T = 20\,\mathrm{min}$, $D = 2\times10^{-3}\,\mathrm{min}^{-1}$ and $D = 3\times10^{-3}\,\mathrm{min}^{-1}$ is caused by evolved cells whose metabolite dynamics oscillates with period $2T$, i.e. twice as large as that of glucose pulse. These cells switch phenotype between imbalanced dynamics during one glucose cycle, and balanced dynamics during the cycle afterwards (Figure S9). Because $B_{\mathrm{p,cov}}$ is defined over an equal number of ON and OFF phases (see *Model and methods*), balancedness of a cell with switching phenotype depends on how many balanced and imbalanced cycles the cell went through and therefore is highly variable. Other high $s(B_{\mathrm{p,cov}})$ values are caused by a few similar genotypes surviving at the end of the simulation due to their similar fitness (akin to the situation in Figure 4, $T = 120\,\mathrm{min}$).

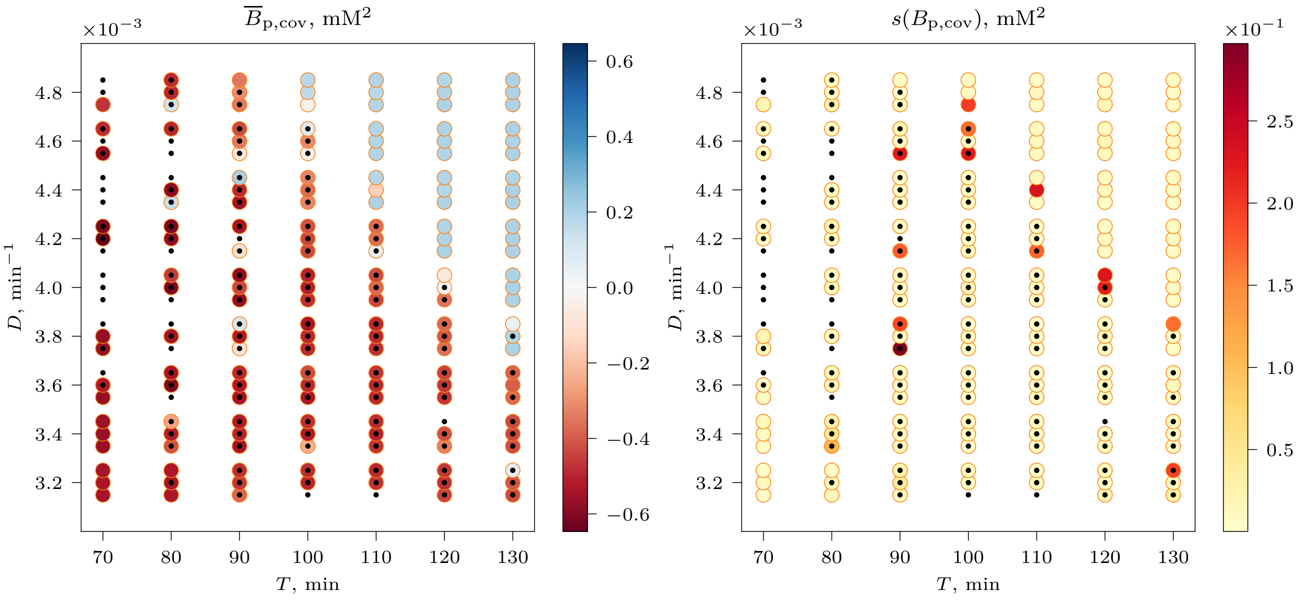


**Figure S6.** Overview of the average phenotypic balancedness $\overline{B}_{\mathrm{p,cov}}$ and its standard deviation $s(B_{\mathrm{p,cov}})$ of evolved tracked cells (colorbar) at the end of an evolutionary simulation ($t_{\mathrm{e}}$) with constant $T_{\mathrm{off}}$ for intermediate values of glucose pulse period $T$ and chemostat flow rate $D$ values. In this range, dimorphism and catastrophic dynamics in the population have often been observed. A group of three or less circles represents three replicate simulations for the same pair of $T$ and $D$ values (i.e., circles have been displaced in the vertical direction for the purpose of visualization; a $D$ value of the group is at the closest tick mark on the vertical axis). A circle in the plot is absent when the population did not survive to the simulation end, or the population size $N(t_{\mathrm{e}}) < 1000$. Black dots indicate simulations where catastrophic dynamics has been observed (see Section *Evolution of increased imbalancedness...*). A black dot without a circle indicates that the population has been wiped out by a catastrophe. Points of high variation in balancedness indicate stable dimorphism in the population, i.e. where both ICs and BCs stably coexist, except for cells at $T = 90\,\mathrm{min}$, $D = 3.8 \times 10^{-3}\,\mathrm{min}^{-1}$ (lower circle) that show phenotype switching (see Figure S5), and cells at $T = 80\,\mathrm{min}$, $D = 3.4 \times 10^{-3}\,\mathrm{min}^{-1}$, where two types of BCs stably coexist.

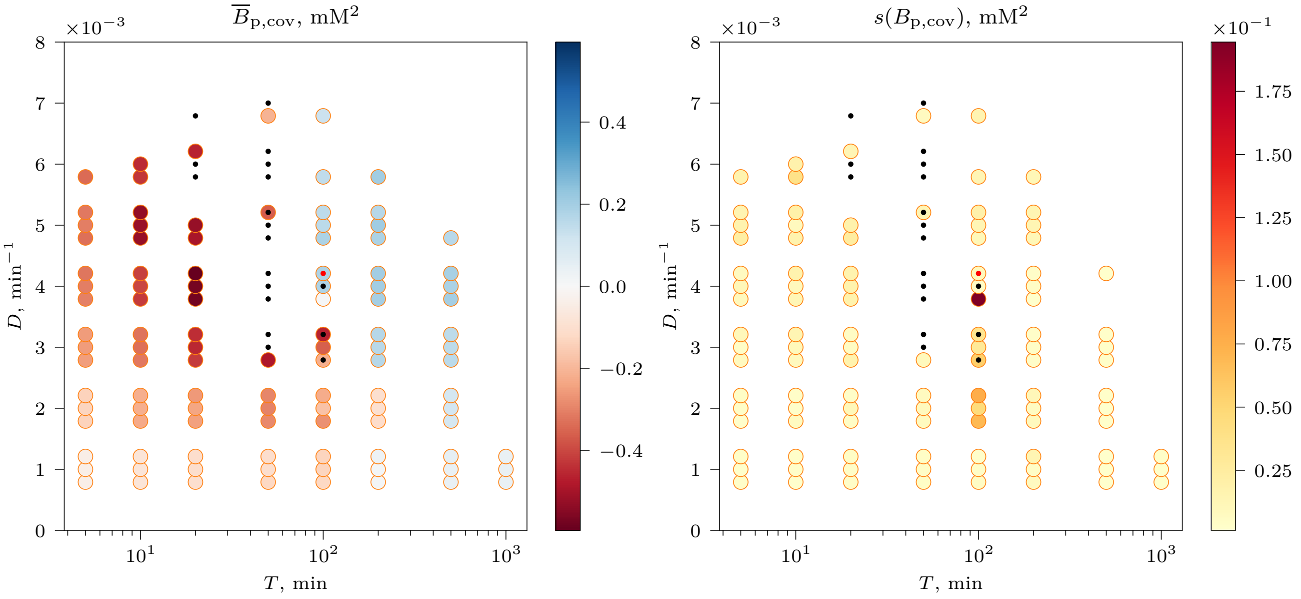


**Figure S7.** Overview of the average phenotypic balancedness $\overline{B}_{\mathrm{p,cov}}$ and its standard deviation $s(B_{\mathrm{p,cov}})$ of evolved tracked cells (colorbar) at the end of an evolutionary simulation ($t_{\mathrm{e}}$) with varying $T_{\mathrm{off}}$, $\mathrm{CV}(T_{\mathrm{off}}) = 5\,\%$, for a wide range of glucose pulse period $T$ and chemostat flow rate $D$ values. A group of three or less circles represents three replicate simulations for the same pair of $T$ and $D$ values (i.e., circles have been displaced in the vertical direction for the purpose of visualization; a $D$ value of the group is at the closest tick mark on the vertical axis). A circle in the plot is absent when the population did not survive to the simulation end, or the population size $N(t_{\mathrm{e}}) < 1000$. Black dots indicate simulations where catastrophic dynamics has been observed (see Section *Evolution of increased imbalancedness...*). One point of high variation in balancedness of cells at $T = 100\,\mathrm{min}$, $D = 4 \times 10^{-3}\,\mathrm{min}^{-1}$ (dark red) is a result of dimorphism in the population, i.e. where both ICs and BCs stably coexist (Figure 5). A black dot without a circle indicates that the population has been wiped out by a catastrophe. The red dot indicate a simulation where stable dimorphism after $t_{\mathrm{me}}$ has been observed, but ICs have been later wiped out by a catastrophe, leaving only BCs in the population.

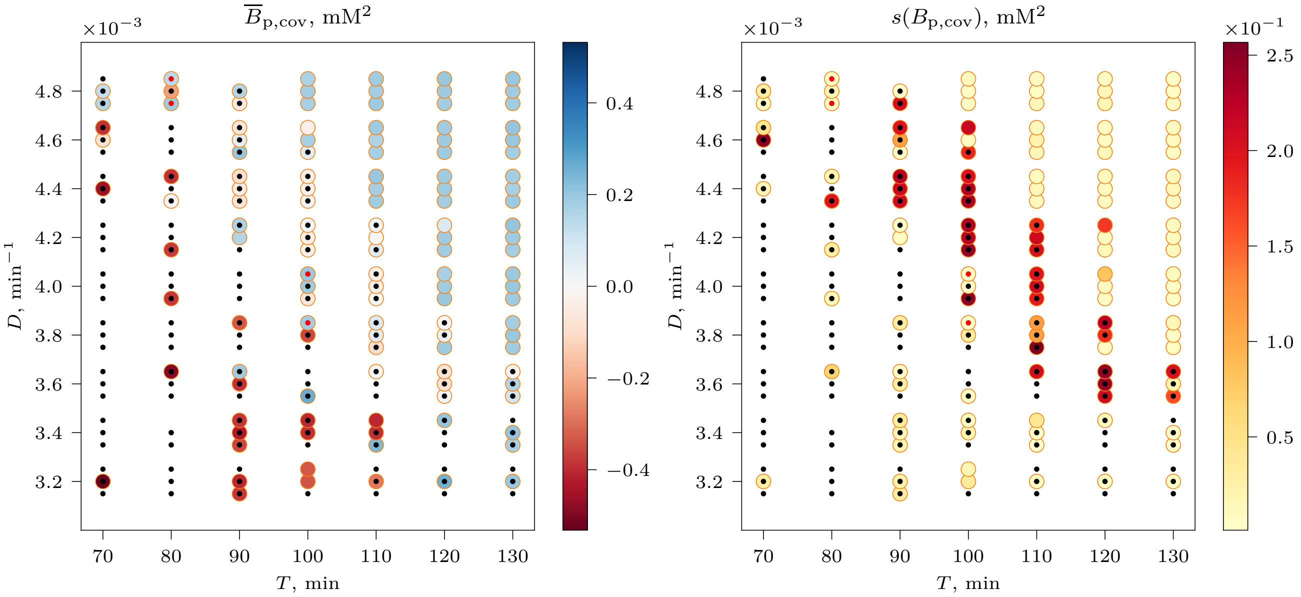


**Figure S8.** Overview of the average phenotypic balancedness $\overline{B}_{\mathrm{p,cov}}$ and its standard deviation $s(B_{\mathrm{p,cov}})$ of evolved tracked cells (colorbar) at the end of an evolutionary simulation ($t_{\mathrm{e}}$) with varying $T_{\mathrm{off}}$, CV($T_{\mathrm{off}}$) = 5 %, for a narrow range of glucose pulse period $T$ and chemostat flow rate $D$ values. In this range, dimorphism and catastrophic dynamics in the population have often been observed. A group of three or less circles represents three replicate simulations for the same pair of $T$ and $D$ values (i.e., circles have been displaced in the vertical direction for the purpose of visualization; a $D$ value of the group is at the closest tick mark on the vertical axis). A circle in the plot is absent when the population did not survive to the simulation end, or the population size $N(t_{\mathrm{e}}) < 1000$. Black dots indicate simulations where catastrophic dynamics has been observed (see Section *Evolution of increased imbalancedness...*). A black dot without a circle indicates that the population has been wiped out by a catastrophe. Points of high variation in balancedness indicate stable dimorphism in the population, i.e. where both ICs and BCs stably coexist. Red dots indicate simulations where stable dimorphism after $t_{\mathrm{me}}$ has been observed, but ICs have been later wiped out by a catastrophe, leaving only BCs in the population. Interestingly, environments with varying $T_{\mathrm{off}}$ are more conducive to forming stable dimorphic populations, compared to the analogous environments with constant $T_{\mathrm{off}}$, where often a monomorphic population of ICs evolves (Figure S6). Thus it appears that variation in $T_{\mathrm{off}}$ is disadvantageous to ICs. This can be explained by the fact that at constant $T_{\mathrm{off}}$, ICs evolve to optimize FBP accumulation so that it is used up just before the next cycle begins. Variation in $T_{\mathrm{off}}$ adds more risk to ICs that the accumulated FBP will not be fully used up during the OFF phase, thus decreasing their fitness and putting them at a disadvantage compared to BCs.

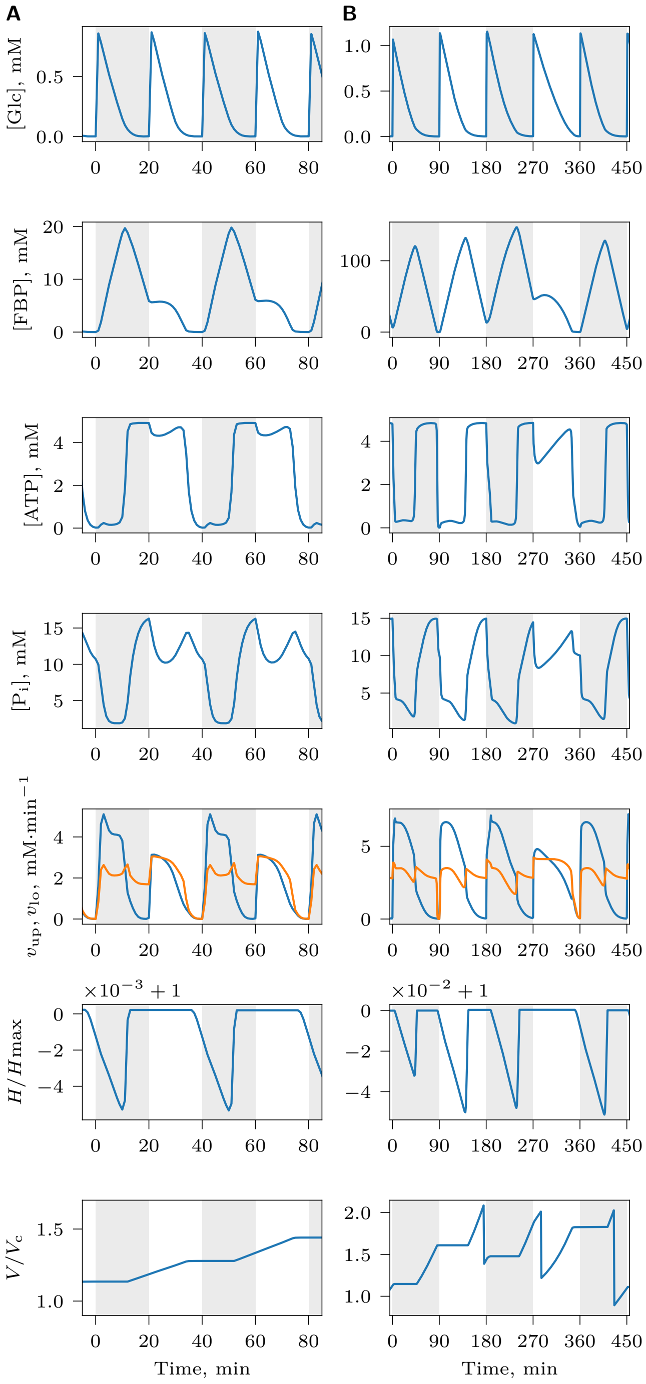


**Figure S9.** Metabolite and growth dynamics of cells with a switching phenotype, evolved in a chemostat at $T = 20\,\mathrm{min}$, constant $T_{\mathrm{off}}$ and $D = 3 \times 10^{-3}\,\mathrm{min}^{-1}$. Cell exhibits both imbalanced and balanced dynamics during different environmental cycles. This occurs because balancedness of glycolysis in the model depends on $[\mathrm{P_i}]$ in the cytosol upon activation with glucose at the beginning of the ON phase (see *Introduction*). (A) Regular phenotype switching. Imbalanced cycle starts with lower $\mathrm{P_i}$, however, due to usage of accumulated FBP, $\mathrm{P_i}$ increases in the cytosol at the end of the cycle, which results in the next cycle being balanced. During the balanced cycle, there is no FBP accumulation in the cytosol, $\mathrm{P_i}$ drops at the end of the cycle, and the next cycle becomes imbalanced again. (B) Irregular phenotype switching.

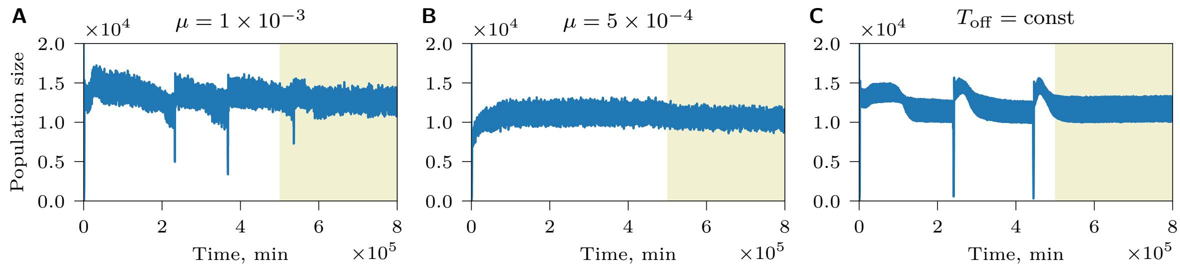


**Figure S10.** Population dynamics in chemostat simulations has less catastrophic events (compared to the dynamics shown in Figure 7A, variable $T_{\text{off}}$, $\overline{T} = 100\,\text{min}$, $\text{CV}(T_{\text{off}}) = 5\,\%$, $D = 4 \times 10^{-3}\,\text{min}^{-1}$ and $\mu = 1 \times 10^{-2}$) when the mutation rate is lower, (A) $\mu = 1 \times 10^{-3}$, (B) $\mu = 5 \times 10^{-4}$, or when (C) $T_{\text{off}}$ is constant. Yellow background indicates the mutation-off segment of the simulation. In populations shown in (A) and (B), dimorphism as in Figure 5 evolves.

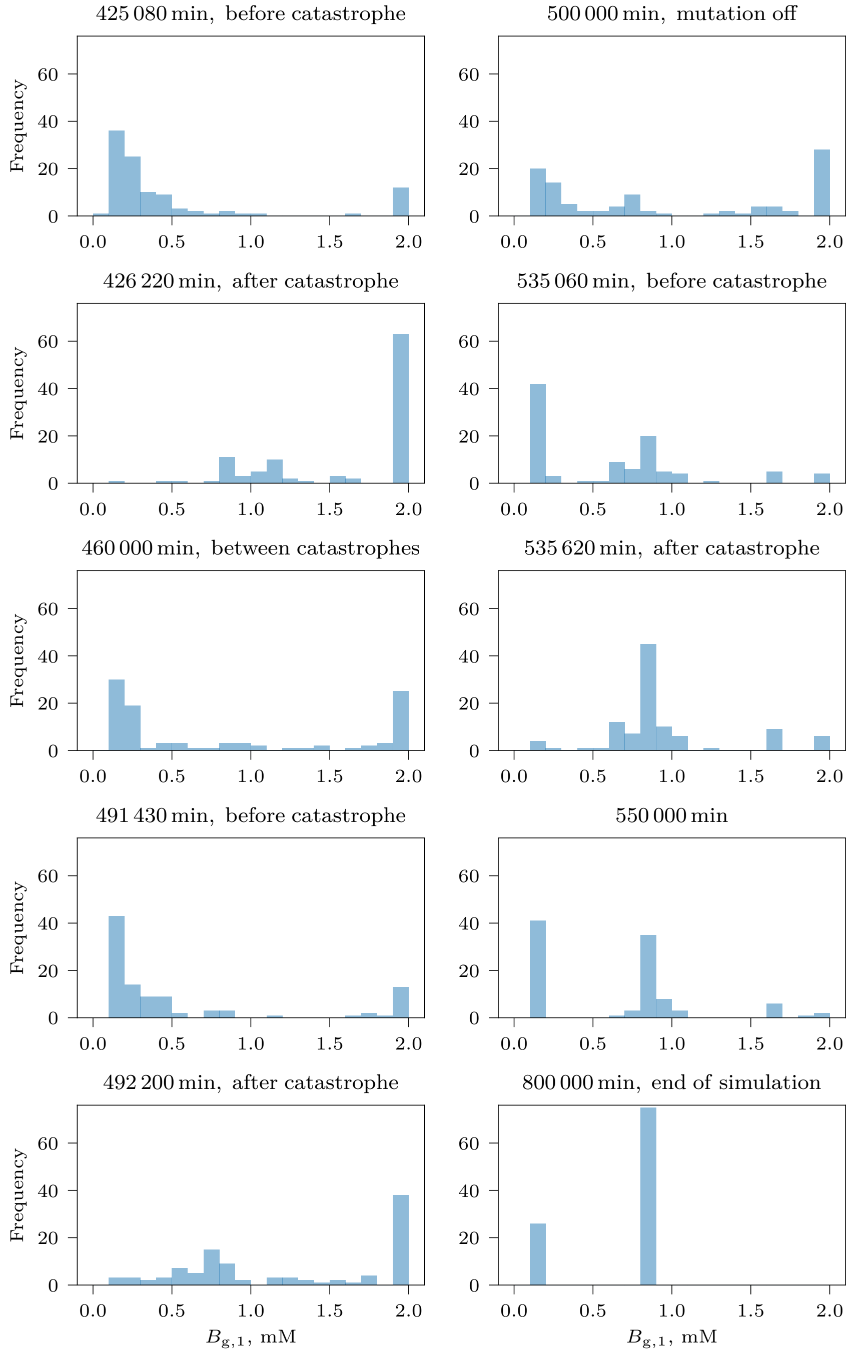


**Figure S11.** Distribution of genotypic balancedness $B_{\text{g},1}$ of tracked cells in time in the simulation shown in Figure 7. During a catastrophe event, the distribution shifts toward more BCs. At the end of the simulation, the population is dimorphic (see Figure 5).

**Supplementary videos**

**Video S1.** Average reproduction rate $r$ of tracked cells during an environmental cycle plotted against their phenotypic balancedness $B_{p,phs}$ during the whole length of a simulation in the NCG scenario with alternating glucose supply, $T = 40$ min (see legend of Figure 4). Yellow background indicates the mutation-off segment of the simulation.

**Video S2.** Average reproduction rate $r$ of tracked cells during an environmental cycle plotted against their phenotypic balancedness $B_{p,phs}$ during the whole length of a simulation in the NCG scenario with alternating glucose supply, $T = 120$ min (see legend of Figure 4). Yellow background indicates the mutation-off segment of the simulation.

**Video S3.** Average reproduction rate $r$ of tracked cells during an environmental cycle plotted against their phenotypic balancedness $B_{p,phs}$ during the whole length of a simulation in the NCG scenario with alternating glucose supply, $T = 200$ min (see legend of Figure 4). Yellow background indicates the mutation-off segment of the simulation.

**Video S4.** Evolution of a dimorphic population. Each dot represents the average reproduction rate $r$ of a tracked cell averaged during an environmental cycle plotted against its phenotypic balancedness $B_{p,cov}$ during a simulation in a chemostat with a variable OFF phase, $\overline{T} = 100$ min, $T_{on} = 1$ min, $\overline{T}_{off} = 99$ min, $CV(T_{off}) = 5\,\%$, and $D = 4 \times 10^{-3}\,\mathrm{min}^{-1}$ (see legend of Figure 5). Yellow background indicates the mutation-off segment of the simulation. At the end of the simulation, subpopulations of BCs and ICs coexist at a stable equilibrium frequency.
